# NEPTUNE: a neuronal cell death pathway prediction tool used to decipher neurodegeneration using single cell RNA-seq datasets

**DOI:** 10.64898/2026.10.31.763288

**Authors:** Ajay Padvi, Alicia A. Brunet, Annie L. Miller, Livia S. Carvalho

## Abstract

In the constantly evolving landscape of cell death mechanisms, much of the literature is dominated by research performed in cancer cells and regenerating cell types. However, neurons differ in that they are non-regenerating and post-mitotic. Therefore, assessing neuronal specific disease biomarkers, including cell death markers, without the bias of non-neuronal or cancerous cells, could provide a more biologically accurate neurodegeneration profile. In this study, we developed the Neuronal cEll death Prediction Tool for Understanding Neurodegenerative Events (NEPTUNE), a user-friendly tool for researchers that bioinformatically identifies neuronal cell death markers from user provided differentially expressed gene lists obtained by single cell RNA sequencing data. We applied NEPTUNE to a publicly available single cell RNA sequencing dataset and analysed the progression of photoreceptor degeneration in the *rd1* mouse model of severe retinitis pigmentosa. The analysis interpreted the dataset in a neuronal-specific manner and showed that late-stage photoreceptor death is not purely apoptotic but possibly engages overlapping cell death programmes. From the analysis, the user is provided with publishable graphs and a list of cell death markers with fold change, type of impact, cell death type and a strength score. Overall, our tool provides markers specific to neurodegeneration, allows mapping to known WikiPathways and KEGG pathways, performs GO and KEGG enrichment analysis and plots the data for further analysis, greatly facilitating the study of neuron specific degeneration. The tool can be found at: https://neptune.rgtcarvalholab.com/.

## Introduction

During the progression of neurodegenerative diseases, it is important to define the underlying mode of cell death to understand how these maladaptive pathways can be therapeutically modulated (Fort, 2023; Green & Llambi, 2015; Krüger & Richter, 2022; Moujalled et al., 2021). With the ever-expanding research into cell death, various modalities have been discovered over time. From the discovery of autophagy in 1963, apoptosis in 1972, and pyroptosis in 2001, the list of defined cell death modalities is progressively increasing (Chen et al., 2024). There are several curated databases such as FerrDb, HAMdb and RCDdb which compile large sets of genes, modulators and their disease associations with various regulated cell deaths that include apoptosis, pyroptosis, necroptosis, autophagy-dependent cell death, entotic cell death, NETotic cell death, parthanatos, mitochondrial permeability transition driven necrosis, immunogenic cell death, lysosome-dependent cell death, and ferroptosis (Wang et al., 2018; X. Wang et al., 2024; Zhou et al., 2026). These databases are comprehensive but can lack specificity because they largely annotate gene-cell death modality relationships without resolving cell type specific effects. Additionally, a substantial amount of the literature used for developing such databases is derived from cancer studies which can confound analysis of non-cancerous cell (Wang et al., 2018; X. Wang et al., 2024; Zhou et al., 2026). It is important to note that multiple cell death regulators exhibit striking context and cell type specificity, even in cancer related cell death. For instance, *CD95/Fas* and its ligand cause classical apoptotic cell death, but has also been shown to have non-apoptotic, pro-survival effects in prometastatic signalling cancer (Fouqué et al., 2014; Haymour et al., 2023; Le Gallo et al., 2017; Seyrek et al., 2024). Likewise, *p53* activation promotes apoptosis in some lymphoid and tumour cells but also induces cell cycle arrest, senescence and survival programmes in other cell types, reflecting a highly context dependent transcriptional network (Aubrey et al., 2018; Lukin et al., 2015).

Neurons, which are largely post-mitotic and lack robust regenerative capacity, show significant vulnerabilities and differences in cell death mechanisms compared to many other cell types (Fricker et al., 2018; Hollville et al., 2019; Stadelmann et al., 2001). For example, neurons are uniquely susceptible to excitotoxic calcium overload via *NMDA/AMPA* receptor mediated Ca²⁺ influx (Guo & Ma, 2021) and *PARP1* mediated parthanatos triggered by bioenergetic collapse under high metabolic demand (Feltes & Alvares, 2024). In retinal photoreceptors, terminal deoxynucleotidyl transferase dUTP nick end labelling (TUNEL) positive cell death proceeds largely in the absence of BAX activation, cytochrome *c* release, and caspase 9 and 3 activity in multiple hereditary retinal degeneration models (Arango-Gonzalez et al., 2014; Brunet et al., 2022; Doonan et al., 2003). Similarly, *YTHDF1*, an inhibitor of ferroptosis in prostate cancer (Y. Wang et al., 2024) can also act as a driver of neuronal ferroptosis (Chang et al., 2025; Liao et al., 2025; Yin et al., 2025). These examples illustrate the divergence between neuronal and canonical cell death pathways, highlighting the need for a neuronal-specific cell death identification tool.

To address these limitations, we have developed the tool named the <u>N</u>euronal c<u>E</u>ll death <u>P</u>rediction <u>T</u>ool for <u>U</u>nderstanding <u>N</u>eurodegenerative <u>E</u>vents (NEPTUNE) which identifies marker genes associated with neuronal cell death from user-supplied differentially expressed genes (DEGs) and their log2 fold change (Log2FC) values derived from single cell RNA sequencing (scRNA-seq) analyses. NEPTUNE not only extracts statistically significant neuron specific death genes but also maps them onto relevant pathways present in WikiPathways and the Kyoto Encyclopedia of Genes and Genomes (KEGG) database, enabling visualisation of their position within canonical signalling cascades. To further analyse the marker genes, Gene Ontology (GO) and KEGG plots can also be created in the tool itself (Agrawal et al., 2024; Kanehisa, 2019; Kanehisa et al., 2025; Kanehisa & Goto, 2000).

To further validate the significance of our database and tool, we applied NEPTUNE to evaluate neuronal rod and cone photoreceptor cell death mechanisms using a published database of scRNA-seq from the *rd1* mouse model of severe inherited retinitis pigmentosa, a degenerative vision loss condition (Chen et al., 2022). Upon using NEPTUNE, we identified a shift in potential cell death mechanisms as photoreceptor degeneration progressed, starting with apoptosis, autophagy and ferroptosis and then shifting to necroptosis and pyroptosis at later stages of degeneration. Thus, NEPTUNE enables researchers to perform neuronal-specific analyses on large transcriptomics datasets to identify potential cell death modalities affecting neurodegeneration in their cell or tissue of choice, visualise significant neurodegeneration markers on WikiPathways and KEGG pathways, perform GO analysis and receive publication ready downloadable interactive plots.

## Materials and Methods

### Finding neuronal cell death specific markers

Our tool focused on extracting the potential markers of regulated cell death types: apoptosis, ferroptosis, necroptosis, autophagy and pyroptosis. To ensure that we were investigating neuron-specific protein or cellular level expression studies, and excluding cancer studies; we used a combination of PubMed Advanced Search Builder and APIs (Application Programming Interfaces) provided by Pubtator3 (Wei et al., 2024). We searched every cell death modality separately and used nomenclature of cell death recommended by the Nomenclature Committee on Cell Death (Galluzzi et al., 2018). Accordingly, we used boolean operators like ‘‘AND’’, ‘‘OR’’ and ‘‘NOT’’. We also used field tags like ‘‘mh’’, ‘‘tiab’’, ‘‘pt’’ and ‘‘sb’’ (Table 1).

**Table 1.** Search strategy used to obtain specific PMIDs.

| Neuron related aliases | (“Neurons”[mh] OR “neuron”[tiab] OR “neurons”[tiab] OR “neuronal”[tiab] OR “Dopaminergic Neurons”[mh] OR “dopaminergic neuron”[tiab] OR “Motor Neurons”[mh] OR “motor neuron”[tiab] OR “Retinal Ganglion Cells”[mh] OR “retinal ganglion cell”[tiab] OR “RGC”[tiab] OR “Photoreceptor Cells”[mh] OR “photoreceptor”[tiab] OR “photoreceptors”[tiab] OR “rod photoreceptor”[tiab] OR “cone photoreceptor”[tiab] OR “rod cell”[tiab] OR “cone cell”[tiab] OR “Retinal Degeneration”[mh] OR “retinal degeneration”[tiab] OR “Retina”[mh] OR “retina”[tiab] OR “retinal”[tiab] OR “macular degeneration”[tiab] OR “age-related macular degeneration”[tiab] OR “AMD”[tiab] OR “Neurodegenerative Diseases”[mh] OR “neurodegeneration”[tiab] OR “neurodegenerative”[tiab] OR “Alzheimer”[tiab] OR “Parkinson”[tiab] OR “Huntington”[tiab] OR “amyotrophic lateral sclerosis”[tiab] OR “ALS”[tiab] OR<br>“hippocampus”[tiab] OR “hippocampal”[tiab] OR “cerebral cortex”[tiab] OR “cortical neuron”[tiab] OR “cerebellum”[tiab] OR “cerebellar”[tiab] OR “substantia nigra”[tiab] OR “spinal cord”[tiab] OR “spinal cord injury”[tiab] OR “optic nerve”[tiab] OR “optic neuropathy”[tiab]) |  |  |  |  |
|---|---|---|---|---|---|
| AND |  |  |  |  |  |
| Expression specificity | ( “Gene Expression Regulation”[mh] OR “gene expression”[tiab] OR “protein expression”[tiab] OR “upregulat*”[tiab] OR “up-regulat*”[tiab] OR “downregulat*”[tiab] OR “down-regulat*”[tiab] OR “overexpress*”[tiab] OR “silenc*”[tiab] OR “knockdown”[tiab] OR “knock-down”[tiab] OR “knockout”[tiab] OR “knock-out”[tiab] OR “knockin”[tiab] OR “knock-in”[tiab] OR “inhibit*”[tiab] OR “activat*”[tiab] OR “promot*”[tiab] OR “deficien*”[tiab] OR “delet*”[tiab] ) |  |  |  |  |
| NOT |  |  |  |  |  |
| Cancer Exclusion | (“cancer”[tiab] OR “tumor”[tiab] OR “tumour”[tiab] OR “carcinoma”[tiab] OR “adenocarcinoma”[tiab] OR “lymphoma”[tiab] OR “leukemia”[tiab] OR “leukaemia”[tiab] OR “melanoma”[tiab] OR “glioblastoma”[tiab] OR “sarcoma”[tiab] OR “hepatocellular”[tiab] OR “colorectal cancer”[tiab] OR “breast cancer”[tiab] OR “lung cancer”[tiab] OR “prostate cancer”[tiab]) |  |  |  |  |
| AND |  |  |  |  |  |
| Cell Death | Apoptosis | Ferroptosis | Necroptosis | Pyroptosis | Autophagy |
|  | (“Apoptosis”[mh] OR “apoptosis”[tiab] OR “apoptotic”[tiab]<br>) | (“Ferroptosis”[mh] OR “ferroptosis”[tiab]) | (“Necroptosis”[mh] OR “necroptosis”[tiab] OR “necroptotic”[tiab] OR “programmed necrosis”[tiab] OR “regulated necrosis”[tiab] OR<br>OR “programmed lytic cell death”[tiab] OR “necrosis”[tiab]<br>OR “necrosis”[mh]<br>) | (“Pyroptosis”[mh] OR “pyroptosis”[tiab] OR “pyroptotic”[tiab] OR “inflammasome-mediated cell death”[tiab] OR “gasdermin-mediated programmed necrosis”[tiab] OR “gasdermin-dependent cell death”[tiab]) | (“Autophagy”[mh] OR “Autophagy/genetics”[mh] OR “Autophagy/metabolism”[mh] OR “autophagy”[tiab] OR “autophagic”[tiab] OR “autophagic cell death”[tiab] OR “type II programmed cell death”[tiab] OR “mitophagy”[tiab] OR “selective autophagy”[tiab] OR “ferritinophagy”[tiab] OR “aggrephagy”[tiab] OR “xenophagy”[tiab] ) |
| AND |  |  |  |  |  |
| (“Immunohistochemistry”[mh] OR “immunohistochemistry”[tiab] OR “IHC”[tiab] OR “immunofluorescence”[tiab] OR “IF staining”[tiab] OR “immunolabeling”[tiab] OR “immunostaining”[tiab] OR “Single-Cell Gene Expression Analysis”[mh] OR<br>“single-cell RNA sequencing”[tiab] OR “scRNA-seq”[tiab] OR “scRNAseq”[tiab] OR “single-cell transcriptomics”[tiab] OR “single cell transcriptomics”[tiab] OR “single-cell sequencing”[tiab] OR “10x Genomics”[tiab] OR “Drop-seq”[tiab] OR “Spatial Transcriptomics”[mh] OR “spatial transcriptomics”[tiab] OR “Visium”[tiab] OR “MERFISH”[tiab] OR “seqFISH”[tiab] OR<br>“spatial gene expression”[tiab] OR “in situ sequencing”[tiab] OR “Blotting, Western”[mh] OR “western blot”[tiab] OR “western blotting”[tiab] OR “immunoblot”[tiab] OR “immunoblotting”[tiab] OR “Proteomics”[mh] OR “proteomics”[tiab] OR “proteomic”[tiab] OR “single-cell proteomics”[tiab] OR “spatial proteomics”[tiab] OR “Immunoprecipitation”[mh] OR “immunoprecipitation”[tiab]) |  |  |  |  |  |
| AND |  |  |  |  |  |
| Only research article inclusion | (“journal article”[pt] OR “preprint”[pt]) |  |  |  |  |
| NOT |  |  |  |  |  |
| Exclusion of other documents | (“review”[pt] OR “systematic review”[pt] OR “meta-analysis”[pt] OR “case reports”[pt] OR “editorial”[pt] OR “letter”[pt] OR “comment”[pt] OR “news”[pt] OR “biography”[pt] OR “congress”[pt] OR “guideline”[pt] OR “practice guideline”[pt] OR “retracted publication”[pt]) |  |  |  |  |
| AND |  |  |  |  |  |
| Filtering for Medline recognised articles | “medline”[sb] |  |  |  |  |

Thus, we retrieved neuron-specific research papers in the form of PubMed IDs (PMIDs) in the order of their relevance with ‘‘best match’’ being at the top (Supplementary dataset S1). These ranked PMIDs we then submitted to the PubTator3 API using inhouse python script which parsed the returned annotations and compiled all the genes mentioned across the retrieved abstracts. Using another python script, we reversed this mapping, indexing each gene to the PMIDs mentioning it. We then retrieved the count for the number of PMIDs that mention each gene (Supplementary dataset S2).

Only genes with more than five PMID mentions were chosen for further analysis. Out of these ranked PMIDs, the top 2-3 most relevant PMIDs will be provided to the user, if any gene is detected from data supplied to the tool. We called these genes ‘‘markers’’. Markers obtained through this analysis were manually curated to further confirm its activity in neurodegeneration. To capture all other potential neuronal cell death markers which were not part of or studied to be part of these cell death types, we exclusively used ‘‘NOT’’ Boolean operator to find research papers which excludes all our curated markers of the above-mentioned five regulated cell death mechanisms and studied their role in neurodegeneration (Table 2). We labelled these markers as ‘‘other neurodegeneration’’ markers. We further classified markers according to their presence in the corresponding KEGG pathway of each cell-death modality (Kanehisa & Goto, 2000). Genes represented in the corresponding KEGG pathway were labelled as ‘‘direct’’. Genes absent from the corresponding pathway but have been reported to play a role in neurodegeneration by the corresponding PMIDs were labelled as ‘‘indirect’’. These labels describe pathway membership and do not define specificity in neuronal cell death. For each marker we assigned regulation (upregulating/downregulating), activity (promoting/inhibiting cell death), prediction (cell death type), found in cell (which type of neuronal cell) and references (PMIDs or DOIs). For the markers which were deemed cell type and context specific, they were classed as ‘‘context-dependent markers’’.

**Table 2.** Search strategy used to obtain other neurodegeneration markers.

| Neuron related aliases | (“Neurons”[mh] OR “neuron”[tiab] OR “neurons”[tiab] OR “neuronal”[tiab] OR “Dopaminergic Neurons”[mh] OR “dopaminergic neuron”[tiab] OR “Motor Neurons”[mh] OR “motor neuron”[tiab] OR “Retinal Ganglion Cells”[mh] OR “retinal ganglion cell”[tiab] OR “RGC”[tiab] OR “Photoreceptor Cells”[mh] OR “photoreceptor”[tiab] OR “photoreceptors”[tiab] OR “rod photoreceptor”[tiab] OR “cone photoreceptor”[tiab] OR “rod cell”[tiab] OR “cone cell”[tiab] OR “Retinal Degeneration”[mh] OR “retinal degeneration”[tiab] OR “Retina”[mh] OR “retina”[tiab] OR “retinal”[tiab] OR “macular degeneration”[tiab] OR “age-related macular degeneration”[tiab] OR “AMD”[tiab] OR “Neurodegenerative Diseases”[mh] OR “neurodegeneration”[tiab] OR “neurodegenerative”[tiab] OR “Alzheimer”[tiab] OR “Parkinson”[tiab] OR “Huntington”[tiab] OR “amyotrophic lateral sclerosis”[tiab] OR “ALS”[tiab] OR<br>“hippocampus”[tiab] OR “hippocampal”[tiab] OR “cerebral cortex”[tiab] OR “cortical neuron”[tiab] OR “cerebellum”[tiab] OR “cerebellar”[tiab] OR “substantia nigra”[tiab] OR “spinal cord”[tiab] OR “spinal cord injury”[tiab] OR “optic nerve”[tiab] OR “optic neuropathy”[tiab]) |  |  |  |  |
|---|---|---|---|---|---|
| AND |  |  |  |  |  |
| Expression specificity | ( “Gene Expression Regulation”[mh] OR “gene expression”[tiab] OR “protein expression”[tiab] OR “upregulat*”[tiab] OR “up-regulat*”[tiab] OR “downregulat*”[tiab] OR “down-regulat*”[tiab] OR “overexpress*”[tiab] OR “silenc*”[tiab] OR “knockdown”[tiab] OR “knock-down”[tiab] OR “knockout”[tiab] OR “knock-out”[tiab] OR “knockin”[tiab] OR “knock-in”[tiab] OR “inhibit*”[tiab] OR “activat*”[tiab] OR “promot*”[tiab] OR “deficien*”[tiab] OR “delet*”[tiab] ) |  |  |  |  |
| NOT |  |  |  |  |  |
| Cancer Exclusion | (“cancer”[tiab] OR “tumor”[tiab] OR “tumour”[tiab] OR “carcinoma”[tiab] OR “adenocarcinoma”[tiab] OR “lymphoma”[tiab] OR “leukemia”[tiab] OR “leukaemia”[tiab] OR “melanoma”[tiab] OR “glioblastoma”[tiab] OR “sarcoma”[tiab] OR “hepatocellular”[tiab] OR “colorectal cancer”[tiab] OR “breast cancer”[tiab] OR “lung cancer”[tiab] OR “prostate cancer”[tiab]) |  |  |  |  |
| AND |  |  |  |  |  |
| Cell Death | Apoptosis | Ferroptosis | Necroptosis | Pyroptosis | Autophagy |
|  | (“Apoptosis”[mh] OR “apoptosis”[tiab] OR “apoptotic”[tiab]<br>) | (“Ferroptosis”[mh] OR “ferroptosis”[tiab]) | (“Necroptosis”[mh] OR “necroptosis”[tiab] OR “necroptotic”[tiab] OR “programmed necrosis”[tiab] OR “regulated necrosis”[tiab] OR<br>OR “programmed lytic cell death”[tiab] OR “necrosis”[tiab]<br>OR “necrosis”[mh]<br>) | (“Pyroptosis”[mh] OR “pyroptosis”[tiab] OR “pyroptotic”[tiab] OR “inflammasome-mediated cell death”[tiab] OR “gasdermin-mediated programmed necrosis”[tiab] OR “gasdermin-dependent cell death”[tiab]) | (“Autophagy”[mh] OR “Autophagy/genetics”[mh] OR “Autophagy/metabolism”[mh] OR “autophagy”[tiab] OR “autophagic”[tiab] OR “autophagic cell death”[tiab] OR “type II programmed cell death”[tiab] OR “mitophagy”[tiab] OR “selective autophagy”[tiab] OR “ferritinophagy”[tiab] OR “aggrephagy”[tiab] OR “xenophagy”[tiab]<br>) |
| AND |  |  |  |  |  |
| (“Immunohistochemistry”[mh] OR “immunohistochemistry”[tiab] OR “IHC”[tiab] OR “immunofluorescence”[tiab] OR “IF staining”[tiab] OR “immunolabeling”[tiab] OR “immunostaining”[tiab] OR “Single-Cell Gene Expression Analysis”[mh] OR<br>“single-cell RNA sequencing”[tiab] OR “scRNA-seq”[tiab] OR “scRNAseq”[tiab] OR “single-cell transcriptomics”[tiab] OR “single cell transcriptomics”[tiab] OR “single-cell sequencing”[tiab] OR “10x Genomics”[tiab] OR “Drop-seq”[tiab] OR “Spatial Transcriptomics”[mh] OR “spatial transcriptomics”[tiab] OR “Visium”[tiab] OR “MERFISH”[tiab] OR “seqFISH”[tiab] OR<br>“spatial gene expression”[tiab] OR “in situ sequencing”[tiab] OR “Blotting, Western”[mh] OR “western blot”[tiab] OR “western blotting”[tiab] OR “immunoblot”[tiab] OR “immunoblotting”[tiab] OR “Proteomics”[mh] OR “proteomics”[tiab] OR “proteomic”[tiab] OR “single-cell proteomics”[tiab] OR “spatial proteomics”[tiab] OR “Immunoprecipitation”[mh] OR “immunoprecipitation”[tiab]) |  |  |  |  |  |
| AND |  |  |  |  |  |
| Only research article inclusion | (“journal article”[pt] OR “preprint”[pt]) |  |  |  |  |
| NOT |  |  |  |  |  |
| Exclusion of other documents | (“review”[pt] OR “systematic review”[pt] OR “meta-analysis”[pt] OR “case reports”[pt] OR “editorial”[pt] OR “letter”[pt] OR “comment”[pt] OR “news”[pt] OR “biography”[pt] OR “congress”[pt] OR “guideline”[pt] OR “practice guideline”[pt] OR “retracted publication”[pt]) |  |  |  |  |
| AND |  |  |  |  |  |
| Getting Medline recognised articles | “medline”[sb] |  |  |  |  |
| NOT |  |  |  |  |  |
| 5 regulated cell death types | (“apoptosis”[tiab] OR “apoptotic”[tiab] OR “Apoptosis”[mh] OR “Ferroptosis”[tiab] OR “ferroptotic”[tiab] OR “Ferroptosis”[mh] OR “Necroptosis”[tiab] OR “necroptotic”[tiab] OR “Necroptosis”[mh] OR “Autophagy”[tiab] OR “autophagic”[tiab] OR “Autophagy”[mh] OR “Pyroptosis”[tiab] OR “pyroptotic”[tiab] OR “Pyroptosis”[mh]) |  |  |  |  |
| NOT |  |  |  |  |  |
| All the curated genes of five regulated cell death types |  |  |  |  |  |

This robust analysis provided us with a set of markers associated with neurodegeneration, along with information about the cell death type, its regulation and the associated PMIDs related to the gene and its cell death function. In this way, we generated an individual marker file for apoptosis, ferroptosis, necroptosis, autophagy, pyroptosis and “other neurodegeneration”. The “other neurodegeneration” pathway was generated to capture other cell death mechanisms that may be involved with the user’s dataset. To update the dataset regularly every six months, we will be utilising the ‘‘create alert’’ option of the PubMed Advanced Search Builder (https://pubmed.ncbi.nlm.nih.gov/advanced/).

### Creating NEPTUNE

To keep the tool self-contained and less dependent of external dependencies, we used ECMAScript 6 (ES6) of JavaScript modules with HTML5 and CSS. We provided ‘‘paste genes’’ and ‘‘file upload’’ alternatives for the user to submit genes and their log2FC values. When the user submits these values, JavaScript reads a positive log2FC as upregulation and a negative log2FC as downregulation. The marker files for all five cell death mechanisms and “other neurodegeneration” were acquired using the built-in Fetch API. Each user-provided gene and its regulation was matched against the marker file using a built-in key-value lookup (Map), keyed on gene symbol and regulation direction (+/−). In this way, genes whose symbol and its direction matched with a marker entry are retained and a “matched marker table” will be created and presented to the user within the tool using the browser’s document object model. To address all the aliases of each gene and exclude ambiguous aliases, we mapped every marker to its NCBI Entrez Gene ID and used that ID to retrieve the official gene symbol and synonyms from the NCBI Gene database using an in-house Python script (Maglott et al., 2011).

To provide comprehensive bar graphs of gene-cell death interaction, we created “fold change scatterplots” and “prediction count bar charts” using HTML5 Canvas 2D API. We used the Graphical Pathway Markup Language, the exchange format of WikiPathways diagrams and subsequent images, to map marker genes detected by the tool and highlight them according to their regulation (Agrawal et al., 2024). If users want to highlight NEPTUNE results of direct markers on KEGG, they our tool has a section to upload the KEGG pathway markup language (KGML) and image of the corresponding cell death type (Kanehisa, 2019; Kanehisa et al., 2025; Kanehisa & Goto, 2000; Wrzodek et al., 2011). We used an inhouse python script for highlighting the markers on KEGG pathways uploaded by the user. We searched WikiPathways for pathways representing each cell death type and selected the one that best matched the framework of gene connections in the corresponding KEGG pathway, comprising of the most direct genes within that pathway (Table 3).

**Table 3.**
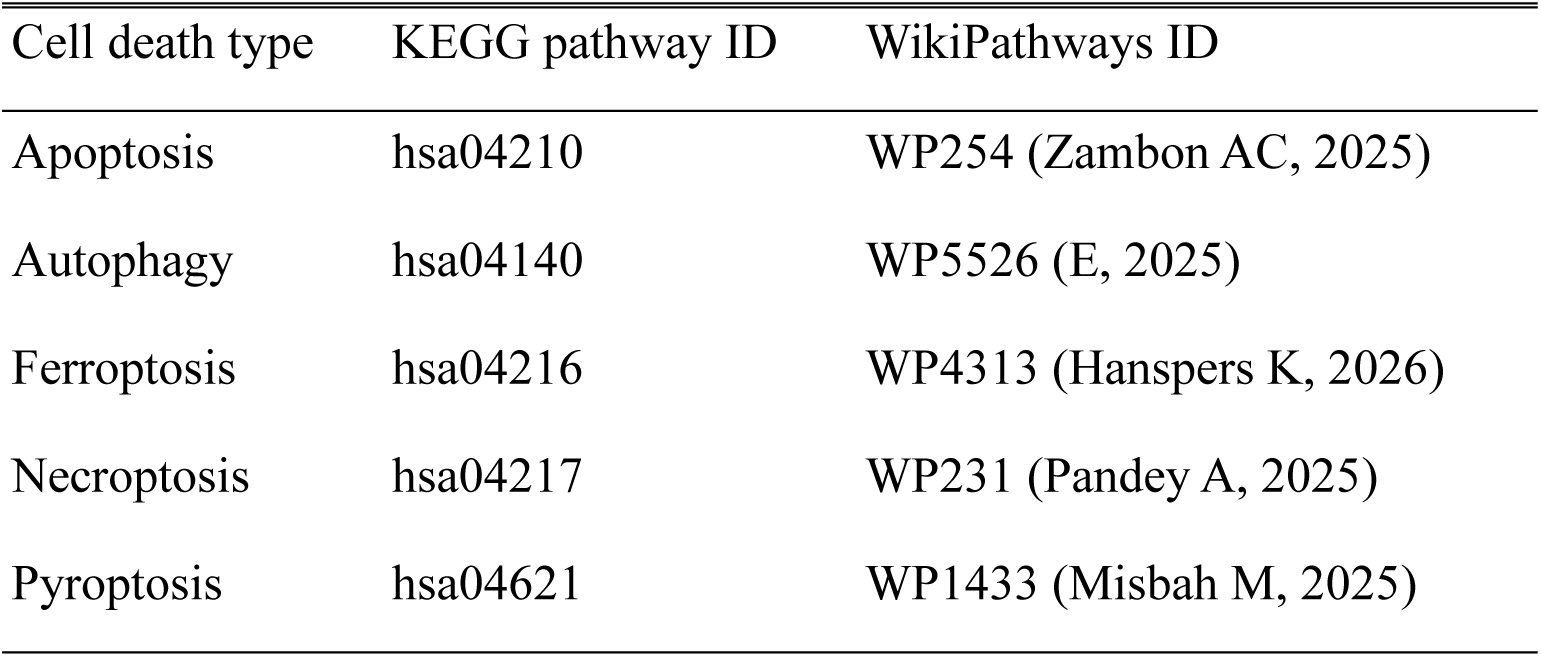
Pathways used for each cell death modality.

Furthermore, to offer users the option to perform over representation analysis, we used REST APIs provided by Enrichr (Chen et al., 2013; Kuleshov et al., 2016; Xie et al., 2021), so that users can utilise the results obtained from NEPTUNE or any DEG dataset they have. We have also provided the option to perform GO analysis (Biological Process, Molecular Function, and Cellular Component) or KEGG pathway analysis using Enrichr libraries (Kuleshov et al., 2016).

### Single Cell RNA sequencing data analysis to evaluate NEPTUNE

To validate NEPTUNE, we used a publicly available dataset of scRNA-seq of retinal cells from the *rd1* mouse model of severe retinitis pigmentosa and wildtype (WT) controls at postnatal days P11, P13 and P17 (Chen et al., 2022). We re-analysed the filtered count matrices deposited by Chen et al. 2022 (GEO accession GSE212183) to obtain differentially expressed genes (DEGs) of *rd1* compared to WT in the rods and cones at P11, P13 and P17.

All analyses were run on the Kaya high-performance computing facility at The University of Western Australia, with jobs submitted through the SLURM workload manager (Infrastructure and Platforms, 2025). The processing was done using the Seurat package (v5) in R 4.5 for normalisation, clustering, annotation and differential expression (Hao et al., 2024). We also used the Matrix package to read the sparse count matrices (Jagan, 2025). We excluded low-quality cells (multiplets or dying cells) based on outlier gene counts and high proportion of mitochondrial reads. We only used cells with 200-5000 detected genes and less than 10% mitochondrial content per cell. Counts were log-normalised, the 2000 most variable genes were identified, and the scaled data was further reduced using principal component analysis before clustering.

Clusters were assigned to retinal cell types using the canonical marker genes reported by the repositor of the dataset (Chen et al., 2022). We used *Rho*, *Nr2e3*, *Nrl*, *Pdc* and *Rp1* for rods; *Opn1sw*, *Opn1mw*, *Pde6h*, *Arr3* and *Gnat2* for cones; *Zfp36l1*, *Dbi*, *Apoe*, *Slc1a3* and *Sparc* for Müller glia; *Pcp2*, *Trpm1*, *Isl1*, *Grm6* and *Trnp1* for bipolar cells; *Meg3*, *C1ql1*, *Snhg11*, *Tfap2b* and *Pcsk1n* for amacrine cells; *Calb1*, *Sept4*, *Tpm3* and *Slc4a3* for horizontal cells; and *Ctsd*, *Ccl4*, *C1qb* and *C1qc* for microglia. After annotating the clusters with these cell specific markers, cell type specific DEGs were extracted using Seurat package on the Kaya HPC facility (Infrastructure and Platforms, 2025). DEGs between *rd1* and WT cells were then filtered within each cell type at each time point using the Wilcoxon rank-sum test, with the thresholds of |avg_log2FC| > 0.58, *p* < 0.05 used by (Chen et al., 2022).

## Results

### Neuronal cell death specific markers

An advanced PubMed search of apoptosis provided 16,217 PMIDs (Table 4). Further, Pubtator3 API extracted 4065 genes mentioned in the respective abstracts of the top 10,000 PMIDs related to apoptosis. After further filtration, where we required each gene to have more than five PMIDs associated and further manual curation, we assigned regulation to 992 markers for apoptosis. This same analysis strategy gave us 262 markers for ferroptosis. For pyroptosis, we extracted 181 markers. Whereas for autophagy, we identified 345 markers. For necroptosis, we identified 205 markers. Excluding the curated markers and cell death they represent, we found 1686 other neurodegeneration markers. Overall, we retrieved 3671 markers from 11,260 genes mentioned in 40,147 neuronal cell death studies of gene or protein expression at a cellular or tissue level.

**Table 4.** Results of our advanced PubMed search using cell death specific search string for apoptosis, ferroptosis, necroptosis, pyroptosis, autophagy, and other neurodegeneration mechanisms. Search string results are number of PMIDs we received after the PubMed advanced search. Genes were retrieved from ranked PMIDs which were ranked based on their relevance. Specific curated markers show number of markers we curated from all the genes received from ranked PMIDs.

| Type of cell death | Search string results | Genes retrieved from cranked PMIDs | Specific curated markers |
|---|---|---|---|
| Apoptosis | 16,217 | 4065 | 992 |
| Ferroptosis | 609 | 617 | 262 |
| Necroptosis | 2703 | 1729 | 205 |
| Pyroptosis | 1,129 | 821 | 181 |
| Autophagy | 2697 | 1896 | 345 |
| Other neurodegeneration markers | 40,147 | 11,260 | 1686 |
|  | 60,805 | 20,388 | 3671 |

*Note: Search was conducted on June 14, 2026.*

### Comparison with FerrDB database

As depicted in Table 5, there was notable overlap with our ferroptotic and apoptotic cell death markers and the corresponding FerrDB database. Out of 262 markers, 137 markers of ferroptosis overlapped with the FerrDB database. Our analysis also provided 114 additional markers which were not mentioned in FerrDB. Similarly, 300 markers of apoptosis were shared between our tool and FerrDB, providing an additional 654 markers which were not present in FerrDB. For necroptosis, 21 out of 205 markers overlapped with FerrDB, while our analysis provided 184 additional markers which were not mentioned in FerrDB. In the case of pyroptosis, 14 out of 181 markers identified by our tool were common with FerrDB, giving 148 additional markers. Likewise, 23 out of 345 autophagy markers overlapped with FerrDB, with our tool contributing 276 markers which were not present in FerrDB. Finally, for the other neurodegeneration markers, 416 out of 1,686 markers overlapped with the FerrDB database when compared across all its regulated cell death mechanisms, while our analysis contributed 1270 additional markers which were not present in FerrDB.

**Table 5.**
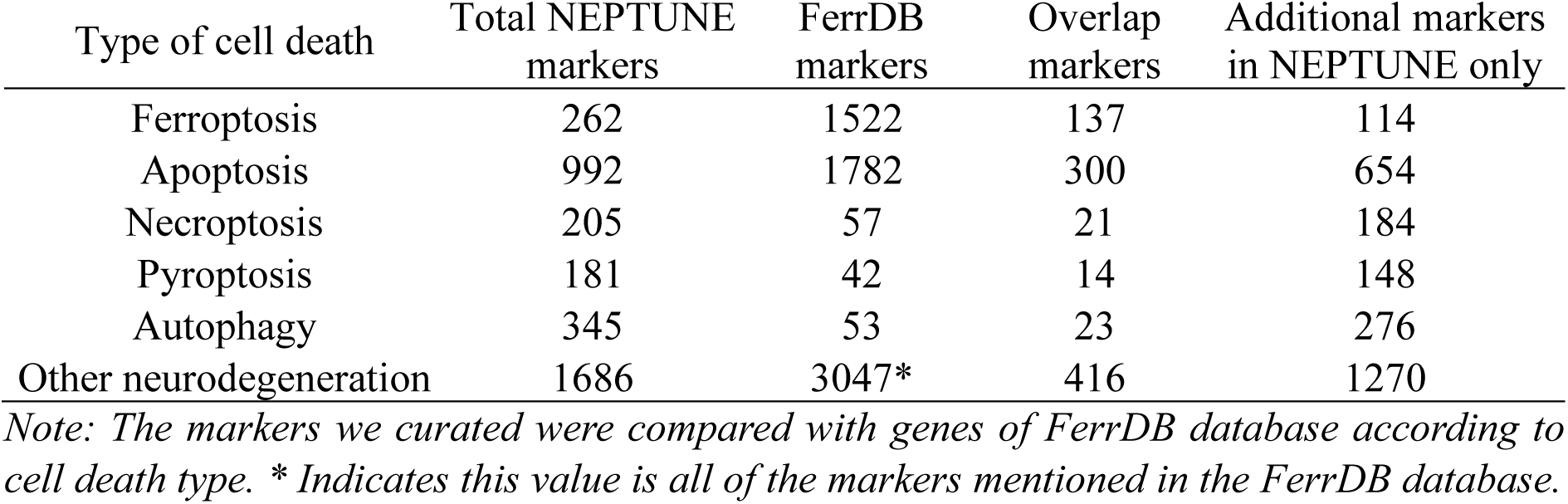
Comparison by type of cell death markers identified using our newly developed NEPTUNE database compared to cell death markers present in the FerrDB dataset.

| Type of cell death | Total NEPTUNE markers | FerrDB markers | Overlap markers | Additional markers in NEPTUNE only |
|---|---|---|---|---|
| Ferroptosis | 262 | 1522 | 137 | 114 |
| Apoptosis | 992 | 1782 | 300 | 654 |
| Necroptosis | 205 | 57 | 21 | 184 |
| Pyroptosis | 181 | 42 | 14 | 148 |
| Autophagy | 345 | 53 | 23 | 276 |
| Other neurodegeneration | 1686 | 3047* | 416 | 1270 |

*Note: The markers we curated were compared with genes of FerrDB database according to cell death type. \* Indicates this value is all of the markers mentioned in the FerrDB database.*

### The NEPTUNE interface

As illustrated in Figure 1, the top of the NEPTUNE webpage provides an overview of the cell death scenario occurring in the neuronal cell, with this data being derived from the DEGs that the user will supply. On the left-hand side of the page, the user has the option to either upload the DEG file or paste the gene list directly. There is also the option to adjust the log2FC threshold as per users’ requirements. On the top right-hand side, we provide options to export the results. The centre of the page consists of an interactive ‘‘log2FC by prediction” plot where each dot showcases a marker and its log2FC values. Additionally, the ‘‘Prediction counts” plot shows the number of markers representing each cell death modality (Figure 1).

**Figure 1.**
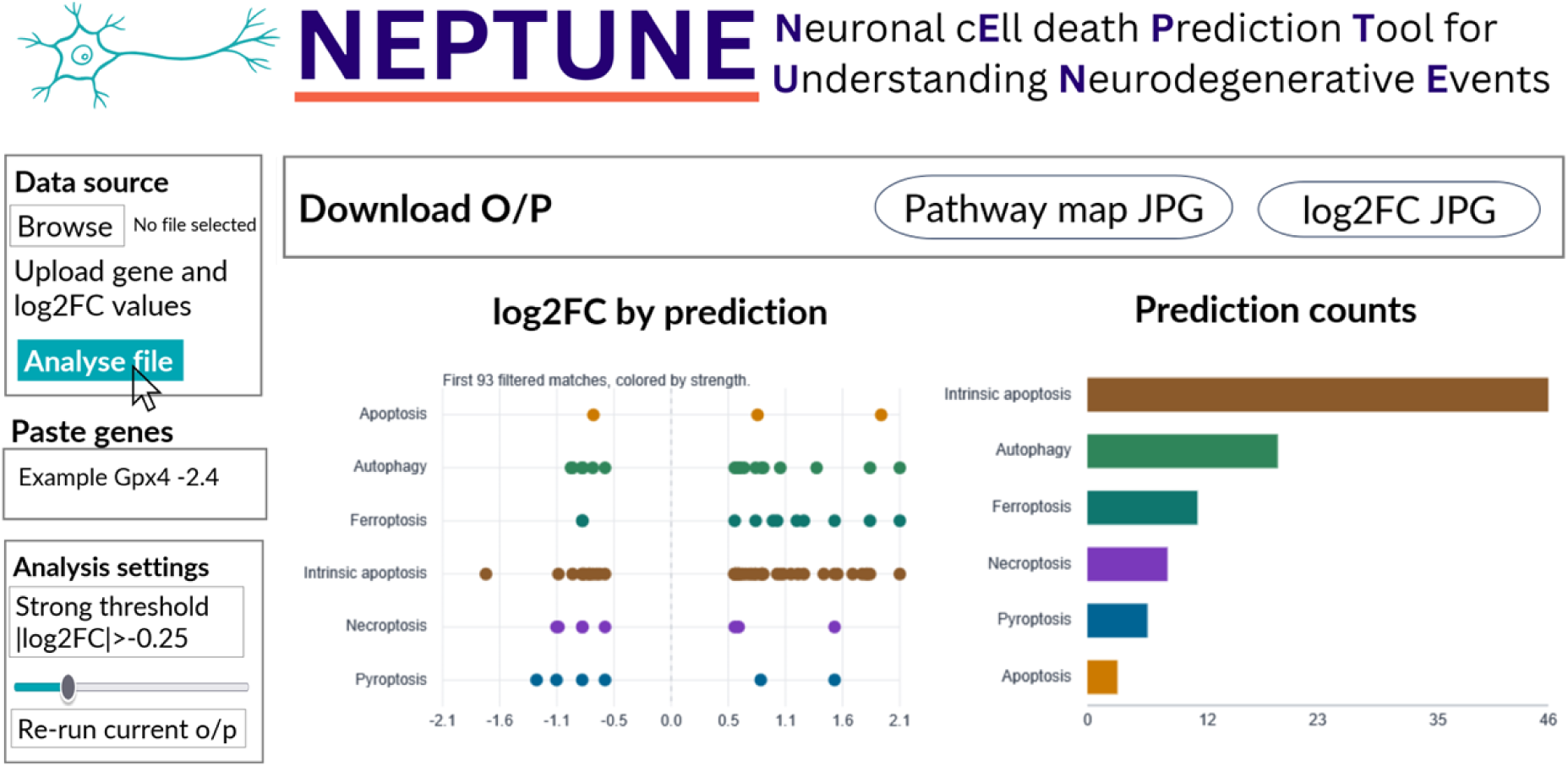
Screenshot showing the NEPTUNE interface where users can input their DEGs and get several types of data. The left side gives the option of ‘‘Data source’’ to upload CSV of gene and its log2FC values. Alternatively, in the bottom left, users can choose to ‘‘Paste genes’’ and the log2FC values of those genes instead of uploading a CSV file. Below the logo, there is an ‘‘Export CSV’’ button to download the NEPTUNE results, a “Pathway map JPG” button to download the WikiPathway diagram associated with their data and ‘‘log2FC plot JPG’’ to download a log2FC by prediction plot.

As shown in Figure 2, the middle of the page shows cell death pathways of WikiPathway highlighting some direct markers of each cell death. Additionally, there is an option to download and upload KGML and KEGG pathway images, to map direct markers found by NEPTUNE on the respective KEGG pathway (Figure 2). The interface also provides an overview of marker counts along with interactive plots of strong and weak markers as per the threshold that is set by the user. The tool also provides the user with an interactive downloadable marker table which showcases a description of each marker, its regulation and neuronal cell type in which it has shown similar regulation (Figure 3). At the end of the page, the tool also gives the option to run GO and KEGG analysis on the dataset that the user has provided, or from the markers provided by the tool. This option allows the user to produce plots and a CSV file as shown in Figure 4.

**Figure 2.**
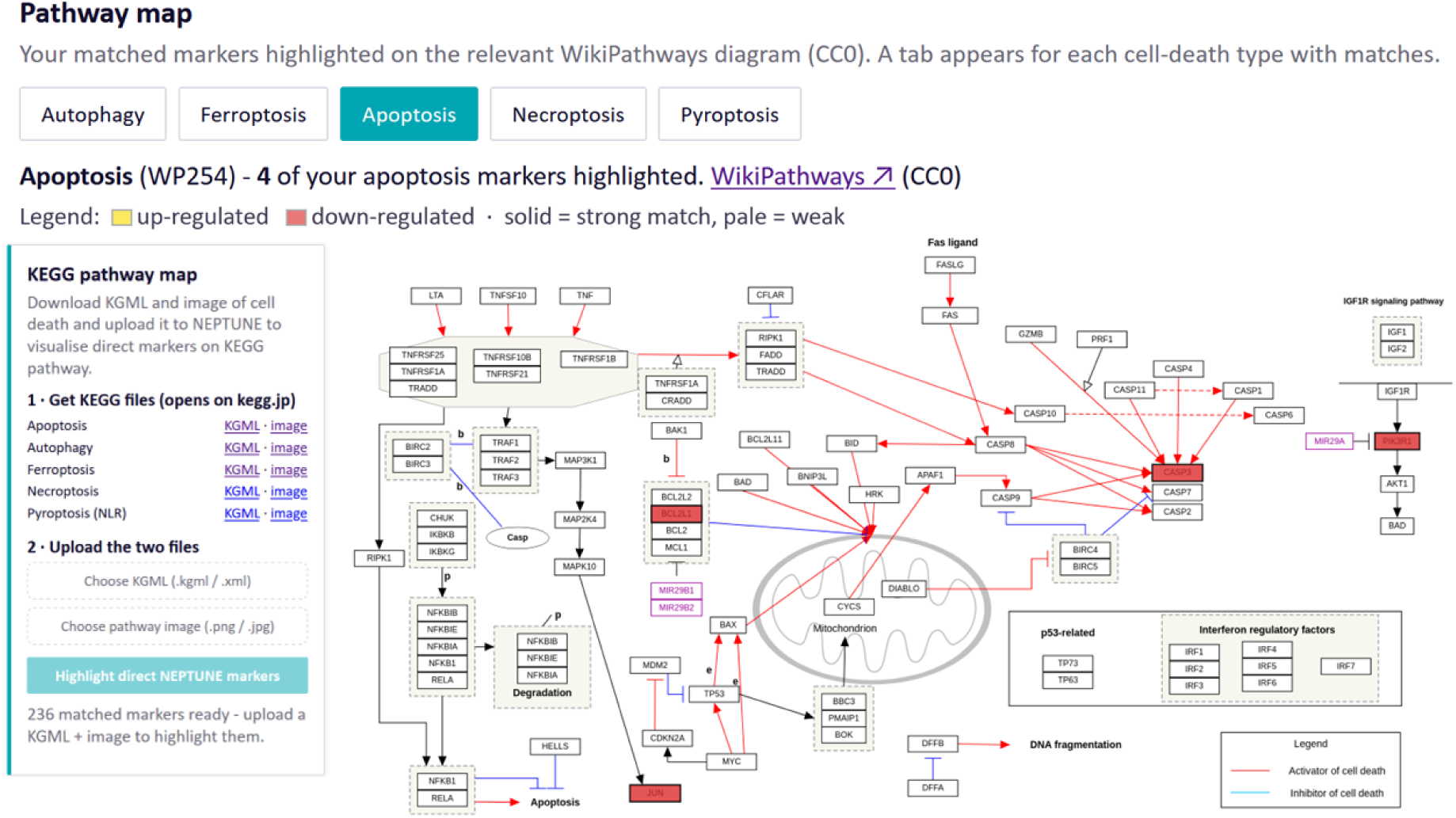
Screenshot of the NEPTUNE interface showing the options to download KGML and KEGG pathway images and upload it back to map the marker genes on the pathway.

**Figure 3.**
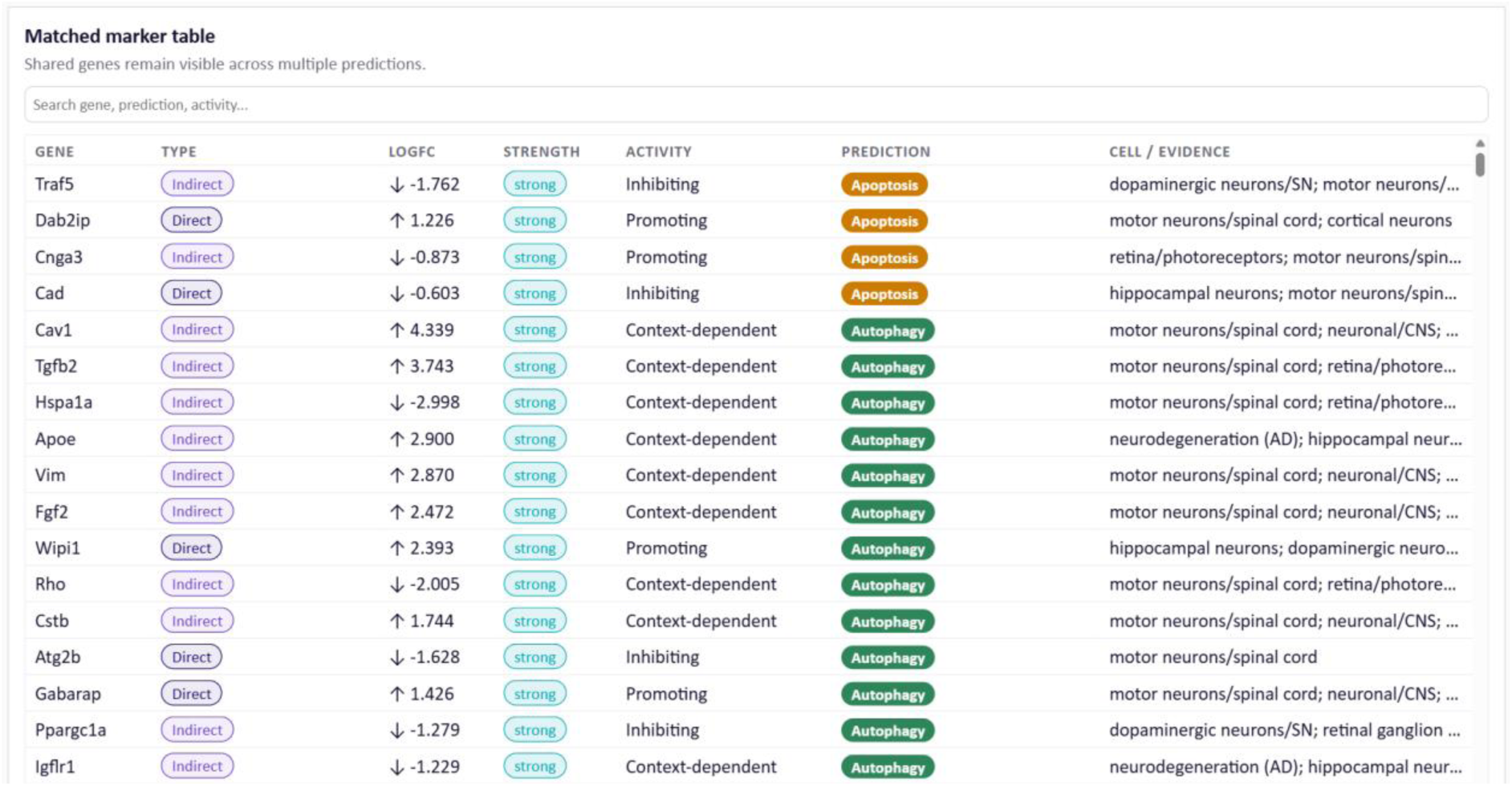
An example of an interactive marker table that is generated for the specific dataset a user provides. ↑ sign shows upregulation and ↓ sign shows down regulation of marker genes detected by NEPTUNE. Each colour shows a different cell death type.

**Figure 4.**
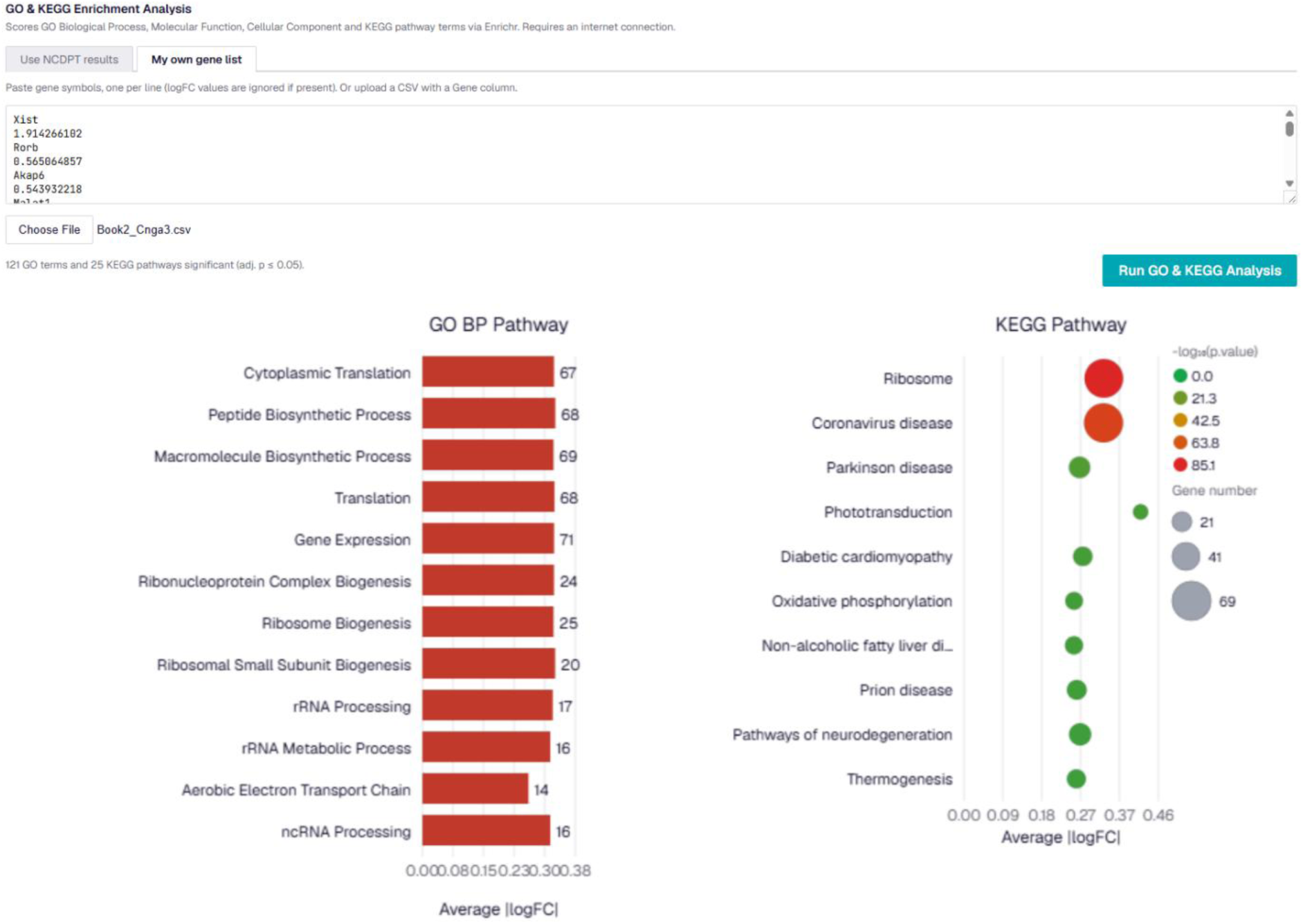
An example of GO and KEGG plots that can be generated by NEPTUNE. GO and KEGG pathway plots were generated using averaged |log2FC values|.

### Evaluation of NEPTUNE using the rd1 Chen et al. 2022 dataset

The clusters formed after analysis of the filtered count matrices were distinct for rods, cones, Müller glia, bipolar cells, amacrine cells and horizontal cells (Supplementary Figures 1-3). We focused only on rod and cone photoreceptor data for downstream analyses, since in *rd1* mice the rods are the mutation-affected cell type that exhibits rapid degeneration, followed by a secondary cone degeneration. This marker-based annotation of clusters provided 5204 rod cells and 92 cone cells in WT retinae at P11, 7924 rods and 172 cones at P13, and 6319 rods and 111 cones at P17 (Supplementary figure 1-3). From *rd1* mice, we obtained 5627 rods and 221 cones at P11, 5803 rods and 495 cones at P13, and 917 rods and 653 cones at P17 ((Supplementary figure 1-3). Cell type and timepoint specific DEGs were then extracted with the Wilcoxon rank-sum test (|avg_log2FC| > 0.58, *p* < 0.05) and we received 598 (P11), 852 (P13) and 1273 (P17) DEGs for rods, and 1167 (P11), 1064 (P13) and 2194 (P17) DEGs for cones (Supplementary Figures 1-3 and Supplementary Table1).

We analysed the DEGs of rod and cone photoreceptors obtained from our analysis of the Chen et al. 2022 dataset using NEPTUNE. In rods, from the 598 DEGs in rd1 versus WT at P11, NEPTUNE detected 19 direct markers of cell death: 23 from 852 DEGs at P13, and 41 from 1273 DEGs at P17 (Figure 5A). In cones, from 1167 DEGs at P11 the tool detected 31 direct markers, this decreased to 21 from 1064 DEGs at P13 and peaked at 57 from 2194 DEGs at P17. For indirect markers, rods showed 74 DEGs at P11, 103 at P13 and 156 at P17 (Figure 5B), while cones showed 113 at P11, 80 at P13 and 206 at P17. Rod cells also showed an increasing number of “other neurodegeneration” markers, with 78 at P11, 110 at P13 and 145 at P17. In cones, a dip at P13 was observed for these markers, with 102 at P11, 77 at P13 and a sharp rise to 230 at P17 (Supplementary dataset S3-S4).

**Figure 5.**
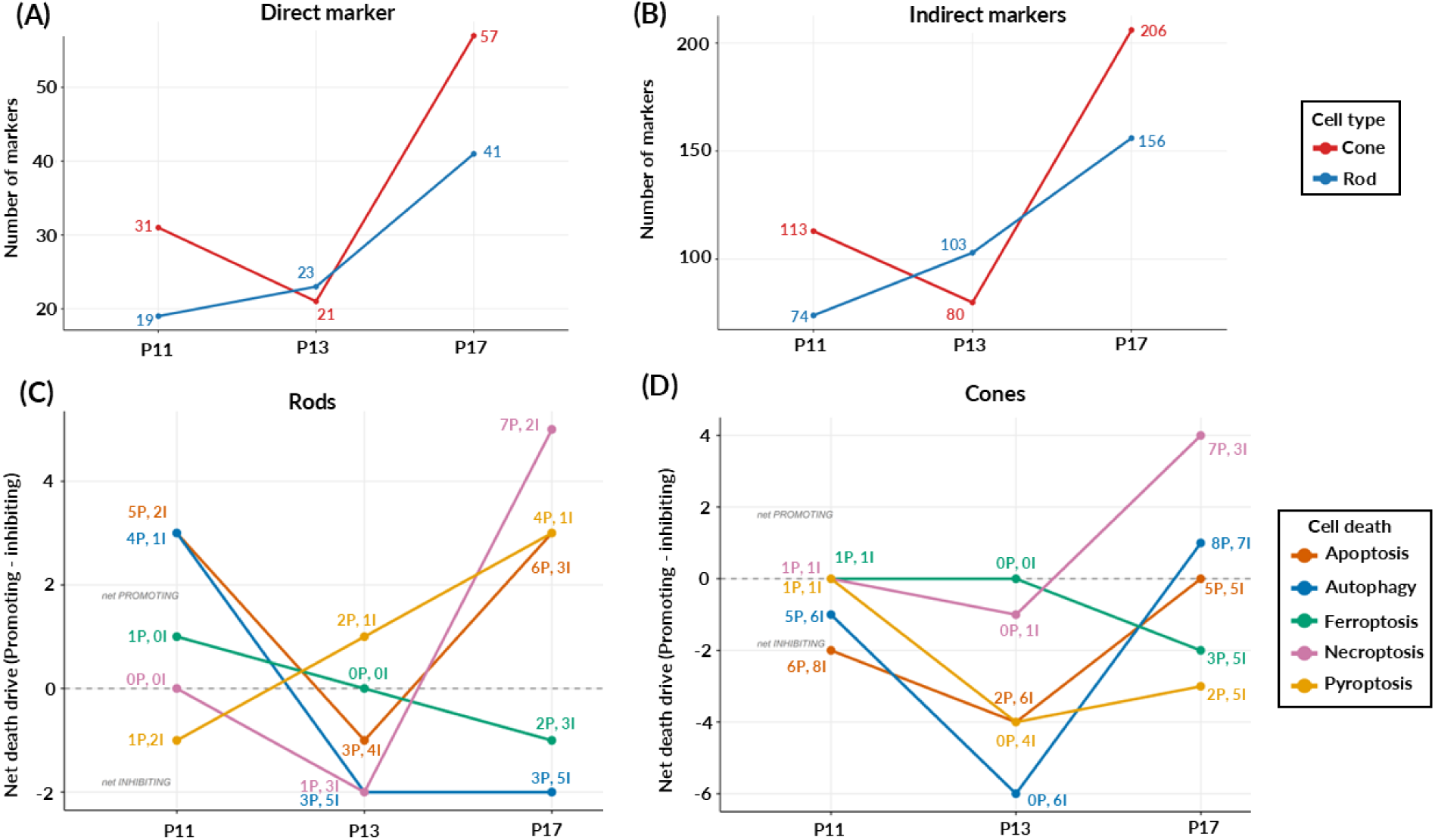
Analysis of direct and indirect markers received from NEPTUNE using DEGs of Chen et al. 2022 dataset of rods and cones in *rd1* mice at P11, P13 and P17. (A) Direct marker counts of rods and cones. (B) Indirect marker counts of rods and cones. Red line indicates number of direct markers for cones at P11, P13 and P17 stages. In (A) and (B), blue line indicates number of direct markers for rods at P11, P13 and P17 stages. (C) Cell death mechanisms shift in degenerating rods over time in the *rd1* mouse model. (D) Cell death mechanisms shift in degenerating cones over time in the *rd1* mouse model. In (C) and (D), the x-axis shows the timepoints (P11, P13, P17) and the y-axis shows net death drive for each cell death mechanism, calculated by subtracting the number of inhibiting direct markers from the number of promoting direct markers. A positive value indicates a net drive toward that death type and a negative value a net suppression. “P” indicates promoting while “I” indicates inhibiting.

### Identifying rod death mechanisms over time using NEPTUNE

Further analysis of DEGs from the Chen et al. 2022 dataset using NEPTUNE was performed on direct cell death markers at P11, P13 and P17 in the rods and cones of the *rd1* mouse model. Figure 5C shows a summary of the direct cell death marker types in rod photoreceptors. At P11 we found five cell death promoting markers of apoptosis, four of autophagy, and one each of ferroptosis and pyroptosis. Two inhibiting markers of apoptosis and pyroptosis were also identified with one of autophagy and none of ferroptosis. At P13, three apoptosis promoting markers were identified, three autophagy and two for pyroptosis. There were no ferroptosis promoting and inhibiting markers at P13. By P17, we identified six promoting markers for apoptosis, three for autophagy, two for ferroptosis, seven for necroptosis and four for pyroptosis. Whereas for inhibiting, we found three markers for apoptosis, five for autophagy, three for ferroptosis, two for necroptosis and one for pyroptosis. Our analysis indicated that overall rod degeneration showed more markers promoting apoptosis and autophagy than inhibiting at P11, suggesting cell death may be mostly driven by these two death modalities at this age. As the degeneration progressed in P13 mice, these cell death types showed an increase in the number of inhibiting markers and a decrease in promoting markers while increasing promoting markers of pyroptosis cell death. The further degeneration seen at P17 showed that cell death may be driven more by necroptosis, pyroptosis and apoptosis with a potential inhibition attempt of other death types such as ferroptosis and autophagy (Figure 5C).

Supplementary figure 4-7 shows the KEGG pathways for the different cell death modalities identified by NEPTUNE in *rd1* rods. The KEGG pathway output diagrams from NEPTUNE also highlighted several upstream regulators of the cell death modalities with similar regulation patterns of promoting or inhibiting the cell death modality in other neuronal cell types, as identified by our curated list of published studies. For example, NEPTUNE identified upregulation of the transcription factor *c-Jun* as a promoter of apoptosis and pyroptosis at P11, however as degeneration progressed at P13, *c-Jun* appeared to be downregulated and labelled as an inhibitor of these same cell death modalities. Similarly, the endoplasmic reticulum (ER) stress sensor kinase *Eif2ak3* (*PERK*) was identified to be inhibiting apoptosis and autophagy at P11 in rods. However, we found other upstream regulators that may have been promoting cell death towards later stages of disease progression. At P13, the mitochondrial checkpoint gene *Bcl2l1* was identified as a promoter of apoptosis, autophagy and pyroptosis. The NF-κB inhibitor *Nfkbia* was flagged as a promoter of apoptosis at P11 but was flipped to inhibiting apoptosis and promoting pyroptosis by P17. The strongest late-stage change at P17 was the deubiquitinase *Tnfaip3* which is also a negative feedback regulator of the NF-κB signalling. *Tnfaip3* was significantly downregulated at P17, being identified as promoting both necroptosis and pyroptosis pathways. At the same stage, the iron storage gene *Fth1* was indicated as an inhibitor of ferroptosis. The results from NEPTUNE also showed research papers showing similar regulation pattern of these markers in the various neuronal cells (Supplementary dataset S4).

### Secondary cone cell death mechanisms over time using NEPTUNE

Cones at P11 were shown to have six apoptosis promoting markers, five autophagy promoting markers and one each of ferroptosis, necroptosis and pyroptosis (Figure 5D). At this stage there were eight apoptosis inhibiting and six autophagy inhibiting markers along with one each for ferroptosis, necroptosis and pyroptosis. By P13, *rd1* cones only showed two apoptosis promoting markers and no other promoting markers of cell death. Instead, NEPTUNE identified several inhibiting markers: six for apoptosis, four for pyroptosis and one for necroptosis (Figure 5D). At P17, cones seemed to shift back towards promoting cell death, with five apoptosis promoting markers, eight for autophagy, seven for ferroptosis, seven for necroptosis and two for pyroptosis. However, there were still several inhibiting markers: five for apoptosis, eight for autophagy, five for ferroptosis, three for necroptosis and five for pyroptosis (Figure 5D). Unlike rods, NEPTUNE identified cones as showing more cell death inhibitory markers than promoting markers at P11 and P13. However, as cone degeneration progresses in the model, the P17 timepoint showed that the number of promoting markers of necroptosis and autophagy significantly increased. Therefore, death drive seems to be led by necroptosis at this age, with potential inhibition attempts through other cell death pathways such as ferroptosis, and autophagy.

Similarly to the rod photoreceptors at P11, the tool suggested that the cones were upregulating the transcription factor *c-Jun.* This was found to be potentially promoting apoptosis and pyroptosis at P11, but its downregulation at P13 could be inhibiting cell death. The ER stress sensor kinase *Eif2ak3* (*Perk*) was also strongly downregulated at P11 as identified in rods and found to potentially inhibit apoptosis and autophagy together with its downstream effector *Eif2a* which showed the same inhibiting pattern. The NF-κB inhibitor *Nfkbia* was flagged as an apoptosis promoter yet was shown to be inhibiting pyroptosis at P11. At P13, the related subunit *Nfkb1* appeared to keep a similar regulation, where apoptosis was promoted while pyroptosis was inhibited. Other upstream regulators showed a shift in gene expression, suggesting they might be more involved in the later stages of cell death. At P17, the tumour suppressor *Trp53* was upregulated and identified as promoting both apoptosis and ferroptosis, while the mitochondrial pore gene *Bax* was upregulated and identified as promoting both apoptosis and necroptosis. The stress kinase *Mapk8* (JNK) was downregulated at P17, potentially driving inhibition of either of apoptosis, necroptosis, pyroptosis and autophagy. The strongest late change at P17 was again the deubiquitinase *Tnfaip3*, which was markedly downregulated and identified as a promoter of both necroptosis and pyroptosis pathways. Additionally, the caspase inhibitor *Xiap* was also indicated to be promoting apoptosis, necroptosis and pyroptosis, consistent with the loss of an apoptosis brake potentially pushing cones toward a more inflammatory death at this age. The markers mentioned above that were identified using NEPTUNE are highlighted in Supplementary Figures 8-11. The results of NEPTUNE also showed research papers showing similar regulation pattern of these markers in the various neuronal cells (Supplementary dataset S4).

### GO and KEGG analysis of photoreceptors using NEPTUNE

In the *rd1* mouse model, rods are primarily affected and are at peak degeneration by P13, while secondary cone degeneration starts at P17 (Chen et al., 2022; Sancho-Pelluz et al., 2008). Taking this into account, we used DEGs obtained from our analysis using NEPTUNE to perform GO and KEGG analysis on rods at P13 and cones at P17 in the *rd1* mouse model. GO analysis showed pathways and DEGs related to vision were enriched, such as detection of visible light, sensory perception of light stimulus and visual perception in rods at P13 (Figure 6A). Whereas KEGG analysis showed the enrichment of pathways including phototransduction and neurodegeneration pathways as well as amyotrophic lateral sclerosis, Huntington disease and Parkinson disease. However, only the phototransduction pathway showed significant enrichment when using adjusted *P*-values (Figure 6B).

**Figure 6.**
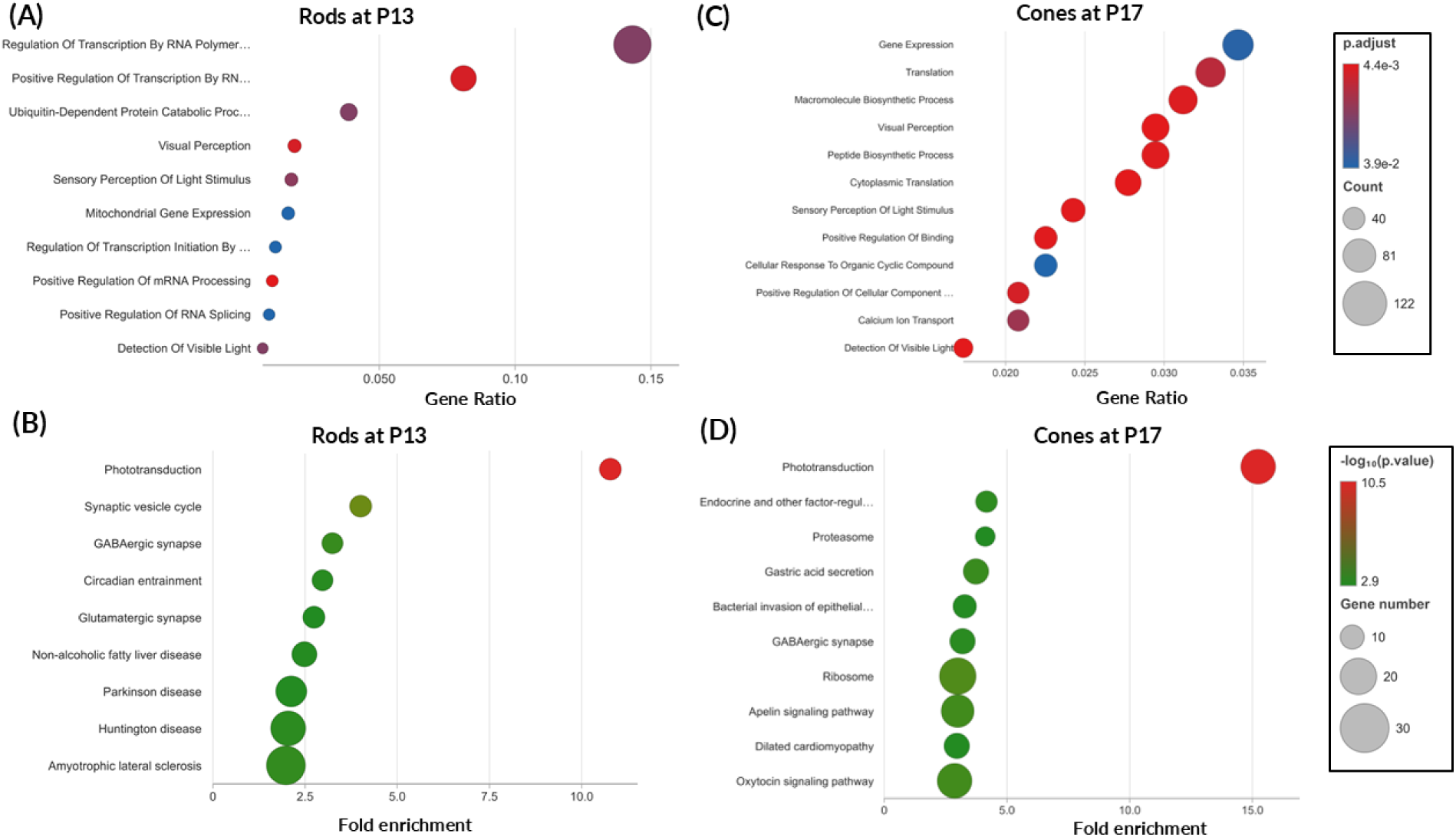
GO and KEGG enrichment analysis performed using NEPTUNE with DEGs of rods and cones received from the Chen et al. 2022 dataset. (A) Rod GO analysis at P13 in the *rd1* mouse. (B) Rod KEGG analysis at P13 in the *rd1* mouse. (C) Cone GO analysis at P17 in the *rd1* mouse model. (D) Cone KEGG analysis at P17 in the *rd1* mouse. For GO enrichment, colour ramp indicates the adjusted *P*-value of each pathway with red being more significant and blue being less. For KEGG enrichment, colour ramp indicates the adjusted *P*-value of each pathway with red being more significant and green being less. Dot size denotes the gene count, the x-axis shows gene ratio, and the y-axis lists the top enriched pathways.

GO analysis in the cone photoreceptors at P17 showed significant enrichment of pathways related to vision, including detection of visible light, sensory perception of light stimulus and visual perception pathways (Figure 6C). Whereas KEGG analysis showed the dysregulation in pathways related to vision such as the phototransduction pathway and cGMP-PKG signalling pathway. Additionally, pathways related to neuronal regulation like GABAergic synapse and oxytocin signalling pathway were also disrupted. However, similar to the rods, only the enrichment of the phototransduction pathway had a significant adjusted *P*-value (Figure 6D).

## Discussion

### NEPTUNE provides an in-silico neuron-specific overview of cell death mechanisms for neurodegenerative diseases

NEPTUNE is a user-friendly platform that provides users with publication ready plots and a holistic overview of the potential degeneration pathways that may be occurring in diseased neuronal cells of interest. Mapping the aberrant cell death markers to previously established WikiPathways and KEGG pathways provides further context and allows easier visualisation of the relevant cell death pathway. The development of this tool was important to remove several limitations that may be encountered when analysing neuronal cells with similar tools available.

Given that NEPTUNE is specifically used for identifying potential cell death pathways relevant to neurons, it was important for us to identify markers for each cell death modality (apoptosis, ferroptosis, pyroptosis autophagy and necroptosis) that have been previously validated. To achieve this, a robust search strategy was developed. After validating NEPTUNE with the Chen et al. 2022 scRNA-seq dataset in *rd1* mice, we found that our search strategy which combines PubMed and Pubtator3 filtered gene specific and neuron specific markers was appropriate. Other than depicting the promotion and inhibition activity of the identified markers, we also provide users context-dependent markers. Such markers may either be promoting or inhibiting cell death depending on the cell type they are expressed in. This information should then be interpreted by the user is per their cell type of interest and related to relevant references provided by NEPTUNE and others.

### Reduction of ambiguity from unrelated pathways

When a user extracts DEGs from their scRNA-seq analysis, the DEG count can be too large to focus on individual cell death genes. Approaches like GO analysis, Gene Set Enrichment analysis and KEGG pathway enrichment analysis allows researchers to connect these DEGs to their biological functions and pathways they play a key role in (Ashburner et al., 2000; Chen et al., 2013; Kuleshov et al., 2016). However, due to a large number of DEGs, untargeted analysis may produce ambiguous results due to pathway crosstalk, gene pleiotropy, and the inclusion of out of context disease categories (Donato et al., 2013; He & Zhang, 2006). As we show in diseased *rd1* cone photoreceptors, alongside canonical neurosensory processes like phototransduction and GABAergic synapse, untargeted analysis indicates significant enrichment for terms such as “bacterial invasion of epithelial cells” and “gastric acid secretion”. These spurious hits may be apparent due to crosstalk of pathogen entry pathways and various degenerative cell death processes such as actin cytoskeletal remodelling, integrin-mediated adhesion, and endocytic trafficking (Stradal & Schelhaas, 2018; Zhang et al., 2026). Since these broad structural categories dominate statistical rankings and top pathway visualisations, representation of specific cell-death related pathways such as apoptosis, necroptosis, or ferroptosis may not be ranked as significantly or sufficiently high compared to other pathways. NEPTUNE addresses this limitation by providing a dedicated, curated view of regulated cell death pathways and their associated genes. Additionally, in the case of scRNA-seq, damaged cells may be lost or filtered out as well as ambient RNA from lysed cells interfering with results, which can distort the profiles of retained cells. Therefore, interpretation of cell degeneration from single cell databases can be skewed by the DEGs of cells that are progressing towards death rather than cells actively undergoing cell death.

A neurodegeneration approach for marker extraction using NEPTUNE not only extracts potential cell death genes from the provided DEGs but also highlights such markers for further investigation. Such markers may reflect an early stress response to cell death and may also explain why multiple intrinsically linked cell death pathways can be enriched at the same time within a specific cellular population. Thus, traditional GO analysis could cause ambiguity as discussed in the previous paragraph, and as we demonstrate in this study. Our tool addresses this ambiguity to an extent, and the previously mentioned limitations by providing neuron specific cell death results.

### Comparison of NEPTUNE with previously established tools confirms its utility

Comparison of NEPTUNE with the previously established FerrDB tool further confirmed the consistency of our approach with previously published databases (Zhou et al., 2026), but with the added novelty of being neuron specific. However, our stringent strategy decreased the number of markers compared to the FerrDb database. For example, we found less similarity with FerrDB’s markers of apoptosis. This could be due to changes in the methodology of identifying apoptosis markers as our analysis was restricted to neuron specific studies. While this may limit the identification of new cell death markers not yet reported in neurons, one of the goals of our search strategy was to avoid a redundancy of markers due to the large number of studies assessing cell death mechanisms in cancer. Our search strategy identified several neuron-specific markers related to these other cell death pathways, making NEPTUNE the only tool currently available with a more comprehensive list of these other cell death modalities. During the development of our tool, there were multiple studies which only showed the role of a gene in neurodegeneration but did not specifically investigate what the cell death type was. In order to account for the presence of these genes linked to neurodegeneration, which could in the future be part of known or new cell death pathways, we also added a “other neurodegeneration markers” category to our list of markers, a unique aspect of our tool. Future updated versions of the tool can revise these genes and change category classification accordingly based on new studies.

### Validation of NEPTUNE with a previously published dataset

To demonstrate the potential of NEPTUNE, we analysed a previously published scRNA-seq dataset of *rd1* mice retinae at the ages of P11, P13 and P17 (Chen et al., 2022). Our tool showed that *rd1* photoreceptor degeneration could potentially progress from pathways linked to apoptosis, autophagy and ferroptosis at P11 to more necroptosis and pyroptosis-like pathways at P17. A similar trend of the cell death progression that our tool identified has been observed previously in multiple mouse models of inherited retinal diseases, where early defects start with autophagy and mitochondrial damage followed by a shift to necroptosis in later disease stages (Newton et al., 2025). However, our study is the first to replicate this finding using a bioinformatic tool. It is also important to note that we focused only on direct markers for this analysis, meaning they were part of corresponding cell death pathways illustrated by KEGG. However, markers being mentioned in KEGG pathways does not necessarily confirm how closely or significantly these direct markers are related to the corresponding cell death pathway. However, NEPTUNE provides the user with the relevant literature to showcase its role in cell death leading to neurodegeneration and the corresponding KEGG pathways which may indicate how these direct markers are leading to cell death. Our study attempted to utilise the connection of genes illustrated by KEGG pathways and studies citing their involvement in particular types of cell death (Dong et al., 2022; Fujikake et al., 2018; Zaidi et al., 2001). Interestingly, some of these direct markers are part of many cell death modalities. For example, the marker *Bcl2l1* was found to be driver of apoptosis, autophagy and pyroptosis in rods at P13 (Tian et al., 2024; Zhong et al., 2022). Therefore, the change in regulation of this marker during disease may lead to one cell death type or several. On the other hand, we also found specific markers such as *Glud1*, *Glul* and *Stat2*, which were found to be promoting only necroptosis in cones at P17 (Atluri et al., 2019; Lim et al., 2020; Tan et al., 2023; Zhao et al., 2024). Therefore, the potential of necroptosis leading to photoreceptor degeneration at P17 cannot be neglected. Similarly, along with shared cell death markers such as *Mapk8*, we also found specific pyroptosis inhibiting markers like *Brcc3, Map3k7* in cones at P13 and *Ikbkg* in cones at P17 (Atluri et al., 2019; Lim et al., 2020; Tan et al., 2023; Zhao et al., 2024). In this way, NEPTUNE was able to interpret the data in the Chen et al. 2022 study from a different perspective and offers a new *in silico* approach for analysing neurodegeneration.

### Considerations when using NEPTUNE

While NEPTUNE provides users with a comprehensive idea about neuron specific markers and their role in neurodegeneration, it is important to note that users should consider what their cell type of interest is as that will alter how they should interpret the data provided by our tool. As such, the user is also advised to further perform molecular validation to confirm that these biological cell death mechanisms are indeed occurring. The neuronal markers identified by NEPTUNE may generally have connotations of what upregulation and downregulation mean and what happens to the cell as a consequence. However, chronic upregulation or downregulation may have the opposite effect. When upregulated, the gene *Hmox1* is considered protective against ferroptosis (de Souza et al., 2026). However, it shows pro-ferroptotic effects when strongly overexpressed (Guo et al., 2024). This phenomenon is also applied in relation to different isoforms of a gene. Length of isoforms may change inhibition or promotion activity in terms of cell death (Boise et al., 1993). For example, long isoform of *Bcl2l1* is antiapoptotic however, short isoform is proapoptotic(Li et al., 2020). Therefore, the user is expected to critically analyse the context-dependent NEPTUNE results within the context of their neuronal cell type, disease or affected area, degeneration stage, and use the citations mentioned in the results, other literature studies and the log2FC value of each marker to guide them. Similarly, as previously discussed, while the automated GO and KEGG analysis that NEPTUNE provides is convenient, it is always important to note that GO analysis may provide users with redundant pathways. As such, users should inspect the csv file of GO results that NEPTUNE provides to further understand the exact context and what pathways are disrupted in the cell. Finally, this tool is currently limited by the fact it only assesses traditional and widely studied cell deaths, specifically apoptosis, ferroptosis, necroptosis, autophagy and pyroptosis. However, other types of death such as parthanatos, sarmoptosis, autosis, autolysis, paraptosis and phagoptosis (Fricker et al., 2018) are important considerations and will be added in future updated versions of the tool. To ensure relevance to the current landscape of cell death mechanisms in neurodegeneration, NEPTUNE will be updated every six to twelve months.

## Conclusions

Neuronal cEll death Prediction Tool for Understanding Neurodegenerative Events (NEPTUNE) which we present in this study may help researchers working in neurodegeneration identify relevant neuron-specific cell death markers which are not biased by other cells such as cancerous cells. The tool not only provides neuron specific markers but also provides users with publication ready plots and identifies potential markers of specific cell death modalities that users can use to guide their further investigations and studies. Validation of the tool against the currently available FerrDB tool and the previously published *rd1* scRNA-seq mouse dataset has shown the utility and efficiency of NEPTUNE in identifying potential markers of cell death, further contributing to our understanding of potential neurodegenerative disease pathways.

## Acknowledgements

The authors acknowledge the use of Kaya, the High-Performance Computing facility at The University of Western Australia, and the technical assistance provided by the UWA IT HPC team (Infrastructure and Platforms, 2025).

## Author contributions

Conceptualisation: AP, AAB, ALM and LSC; methodology: AP; formal analysis: AP, writing (original draft): AP; writing (reviewing and editing): AAB, ALM, LSC; funding: LSC; supervision: AAB, ALM, LSC.

## Data availability

The manuscript includes all data generated or analysed during this study.

