## Supplementary data for "NEPTUNE: a neuronal cell death pathway prediction tool used to decipher neurodegeneration using single cell RNA-seq datasets"

### Supplementary material

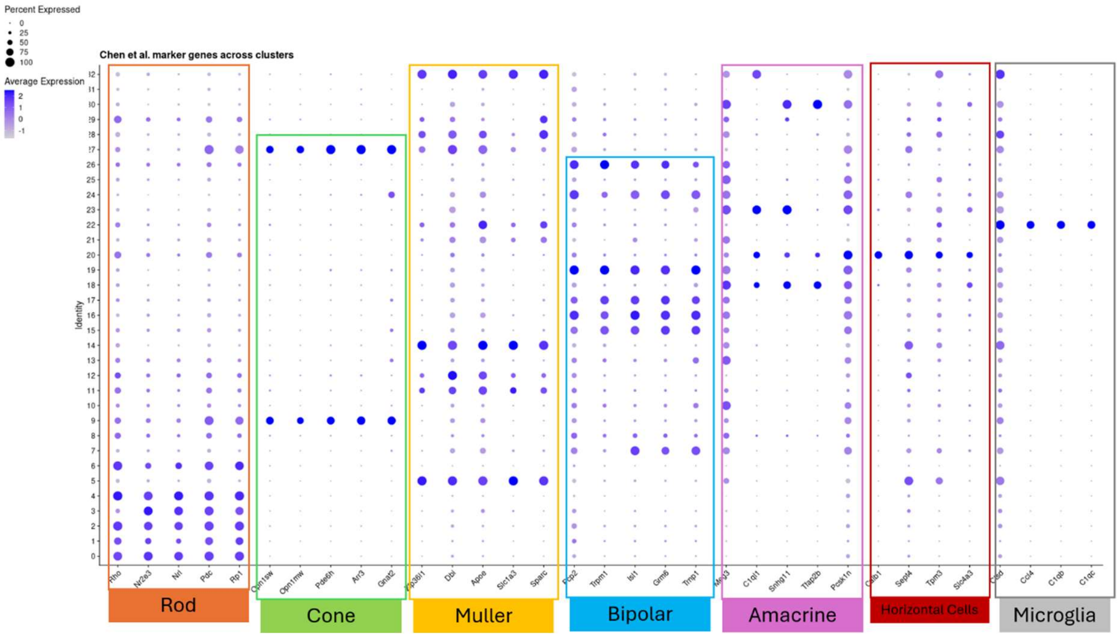


Supplementary Figure 1. Cell clustering using markers identified by the (Chen et al., 2022) dataset that we used to validate NEPTUNE. The DotPlot function of Seurat was used to visualise the expression of canonical retinal marker genes across clusters so that cell-type identity could be confirmed. Genes were plotted on the x-axis and grouped by the cell type they mark; the y-axis denotes the cluster number, with the assigned cell type shown at the bottom. The colour ramp indicates the average expression of each gene within a cell population, and dot size indicates the percentage of cells expressing that gene in that population. Cluster 9 and 27 were annotated as cones. Clusters 5, 11, 12, 14, 28 and 32 were identified as Müller glial cells. Clusters 7, 15, 16, 17, 19, 24 and 26 were annotated to bipolar cells. Clusters 18, 23 and 30 were annotated to be amacrine cells. Further cluster 20 for horizontal cells, cluster 22 microglia and other clusters are left unannotated.

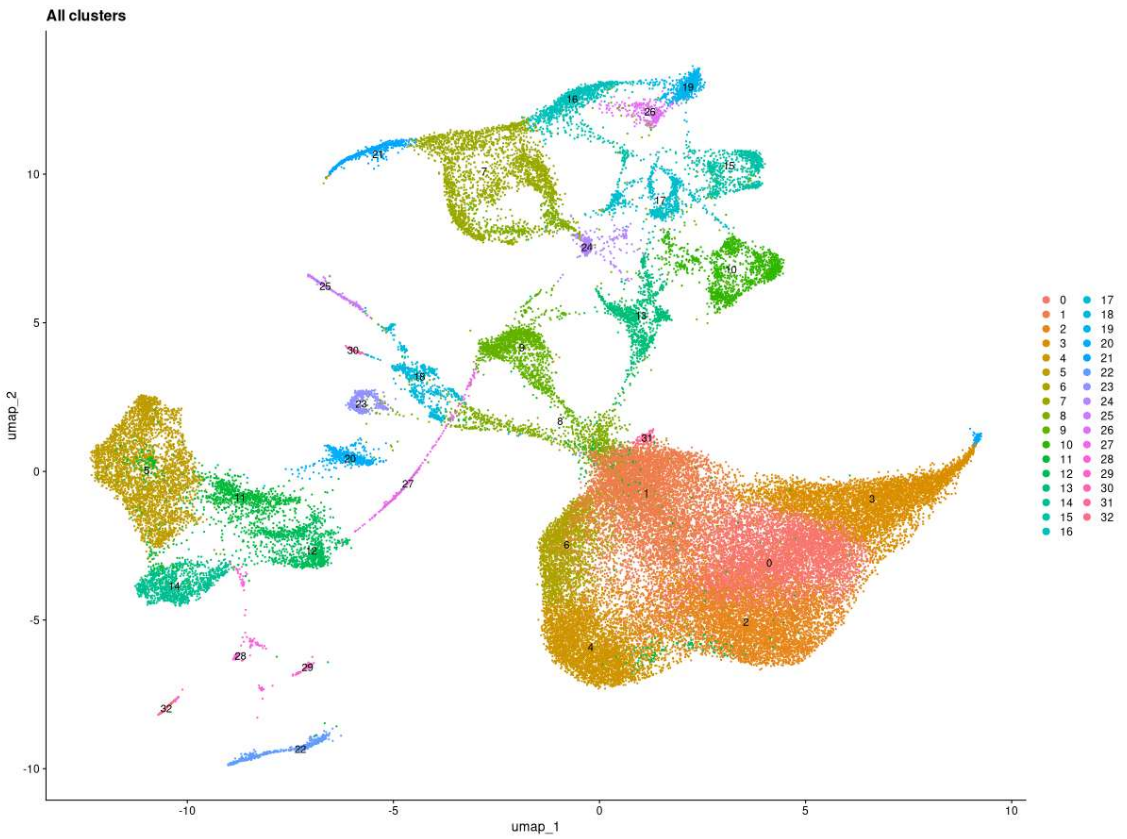


Supplementary Figure 2. Clusters formed after single cell analysis of the WT vs *rd1* mouse retina dataset analysed from (Chen et al., 2022).

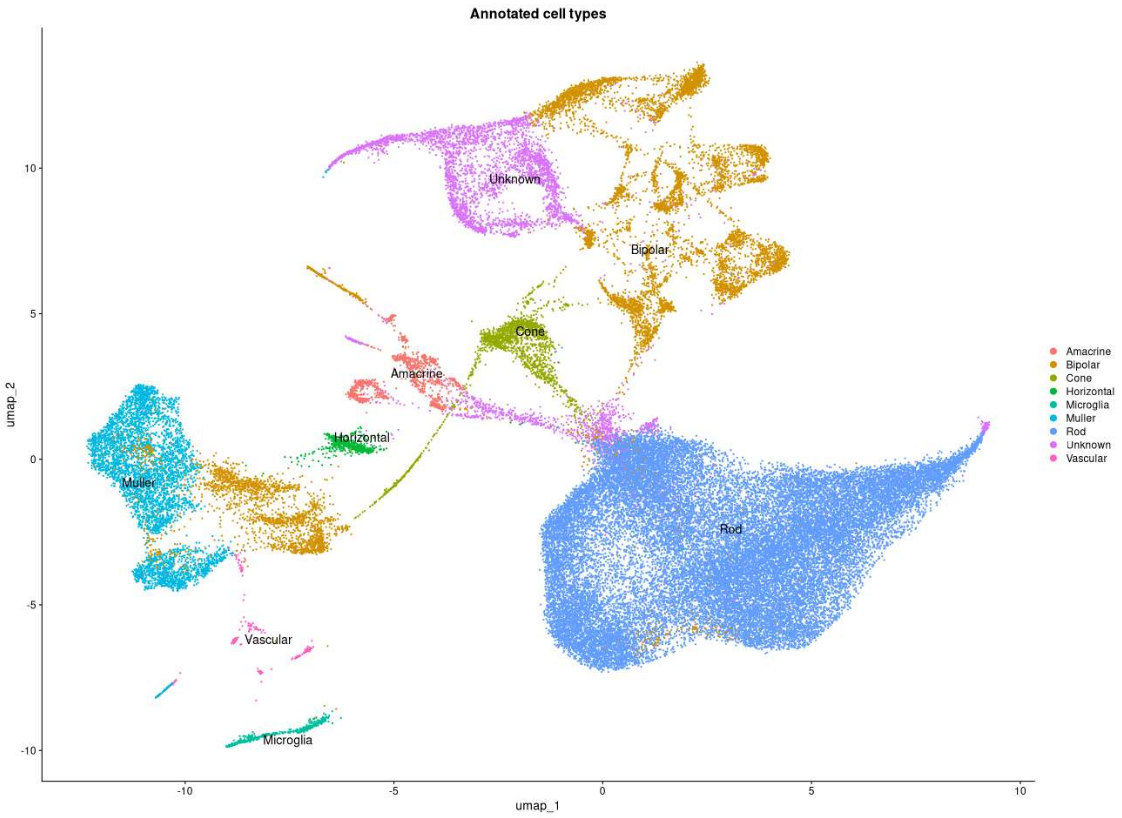


Supplementary Figure 3. Annotated clusters formed after single cell analysis of WT vs *rd1*. Annotated clusters are assigned different colours. The rod cluster is blue while the cone cluster is green.

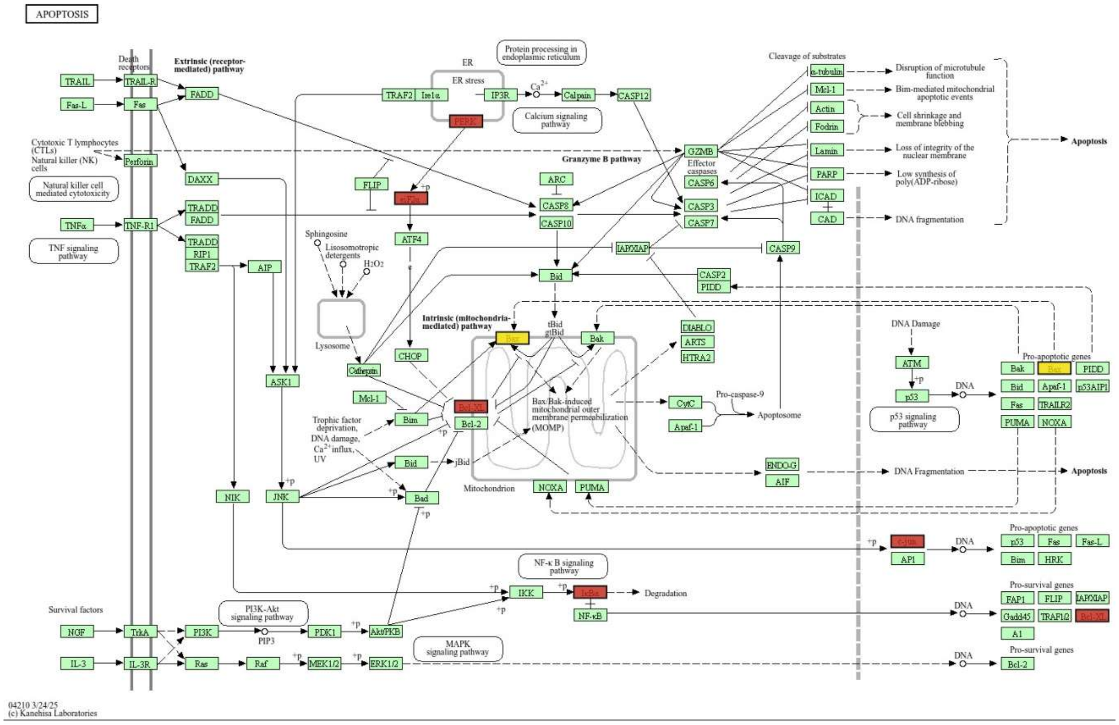


Supplementary Figure 4. NEPTUNE highlighted direct markers of apoptosis across all timepoints during *rd1* rod cell death. Red labels indicate downregulated markers while yellow labels indicate upregulated markers. Pale colours show weak regulation dictated by the |log2FC| values. Green colour indicates all the other genes of the pathway.

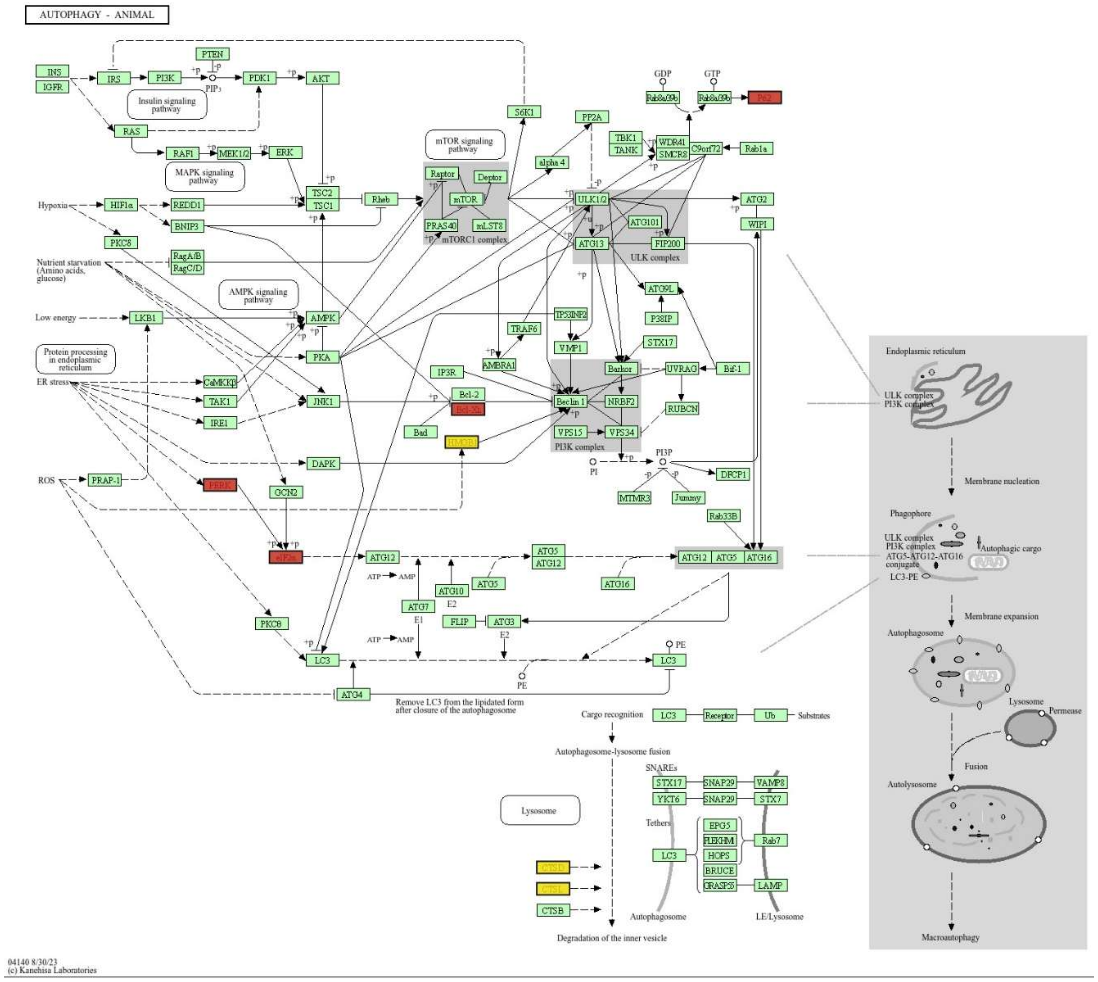


Supplementary Figure 5. NEPTUNE highlighted direct markers of autophagy across all timepoints in *rd1* rod cells. Red labels indicate downregulated markers while yellow labels indicate upregulated markers. Pale colours show weak regulation dictated by the |log2FC| values. Green colour indicates all the other genes of the pathway.

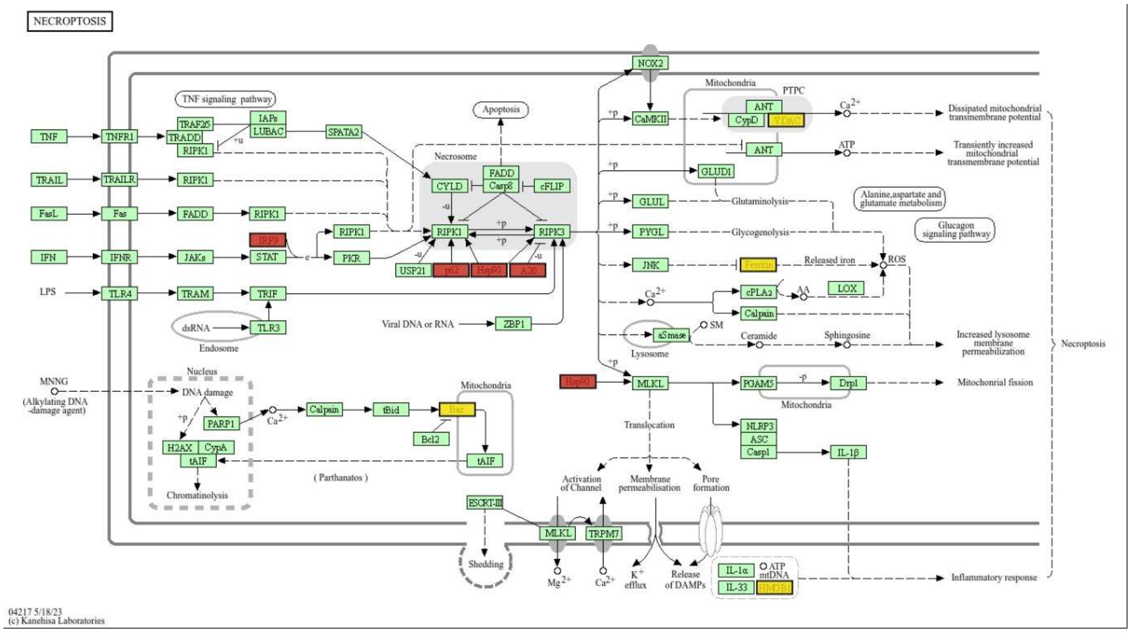


Supplementary Figure 6. NEPTUNE highlighted direct markers of necroptosis across all timepoint in *rd1* rod cells. Red labels indicate downregulated markers while yellow labels indicate upregulated markers. Pale colours show weak regulation dictated by the |log2FC| values. Green colour indicates all the other genes of the pathway.

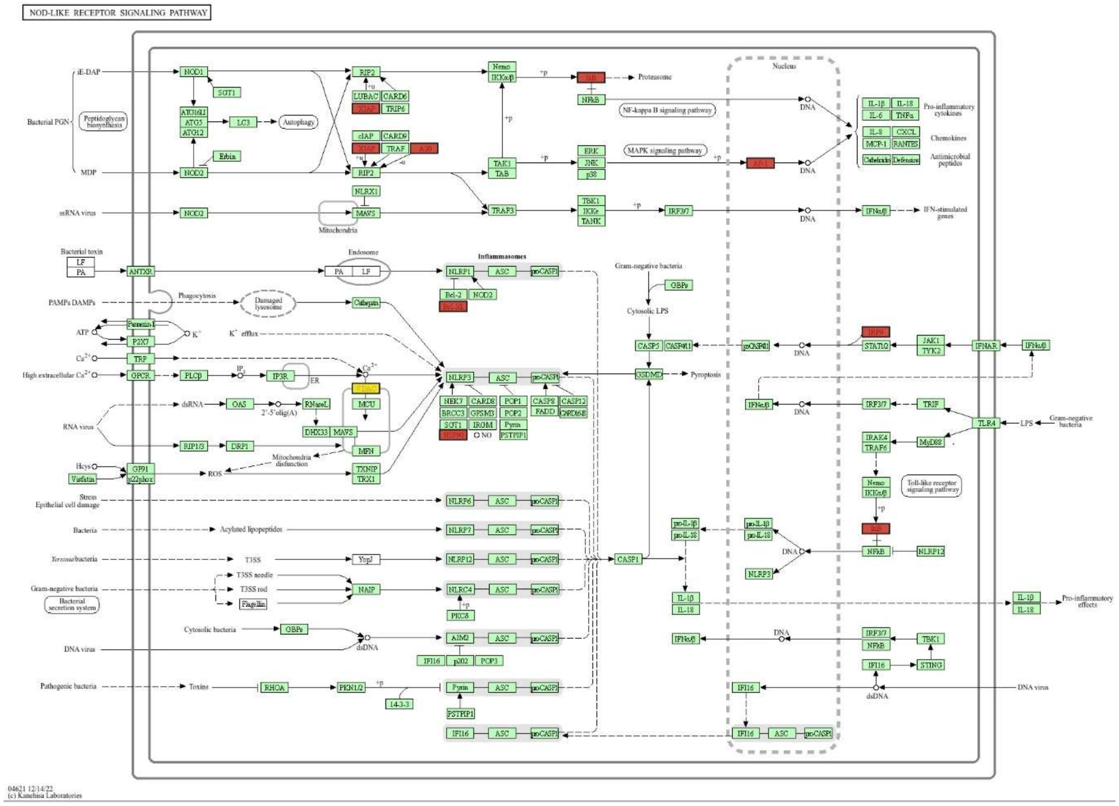


Supplementary Figure 7. NEPTUNE highlighted direct markers of pyroptosis across all timepoint in *rd1* rod cells. Red labels indicate downregulated markers while yellow labels indicate upregulated markers. Pale colours show weak regulation dictated by the |log2FC| values. Green colour indicates all the other genes of the pathway.

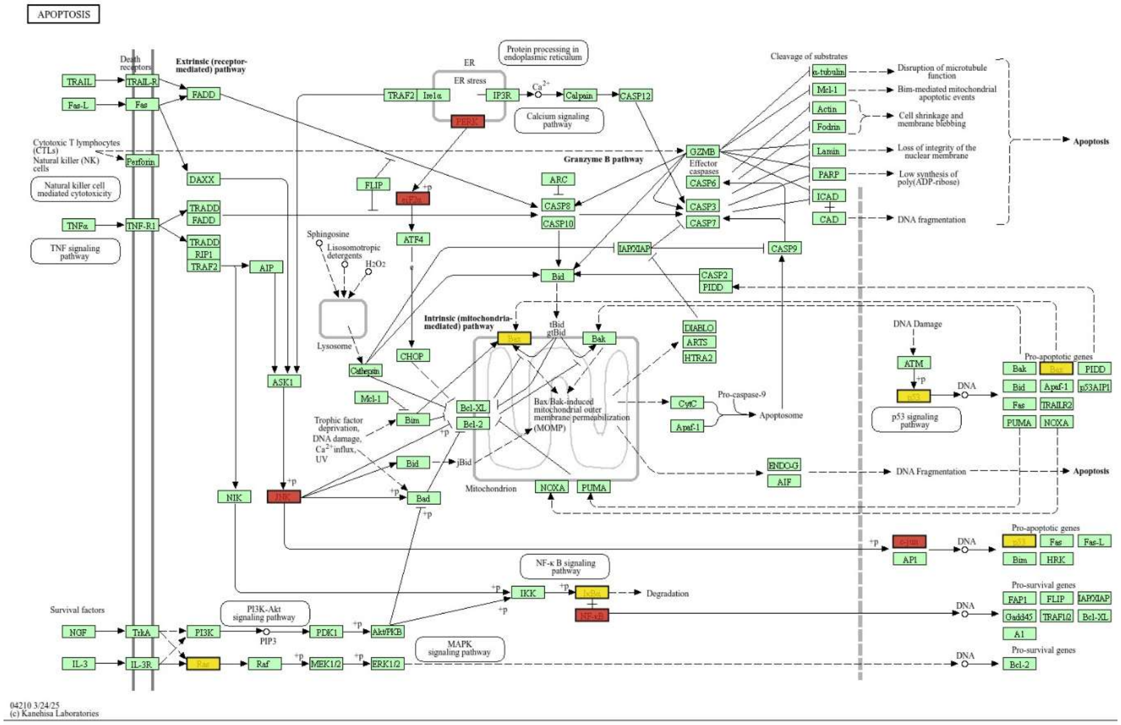


Supplementary Figure 8. NEPTUNE highlighted direct markers of apoptosis across all timepoints in the cones of *rd1* mice. Red labels indicate downregulated markers while yellow labels indicate upregulated markers. Pale colours show weak regulation dictated by the |log2FC values. Green colour indicates all the other genes of the pathway.

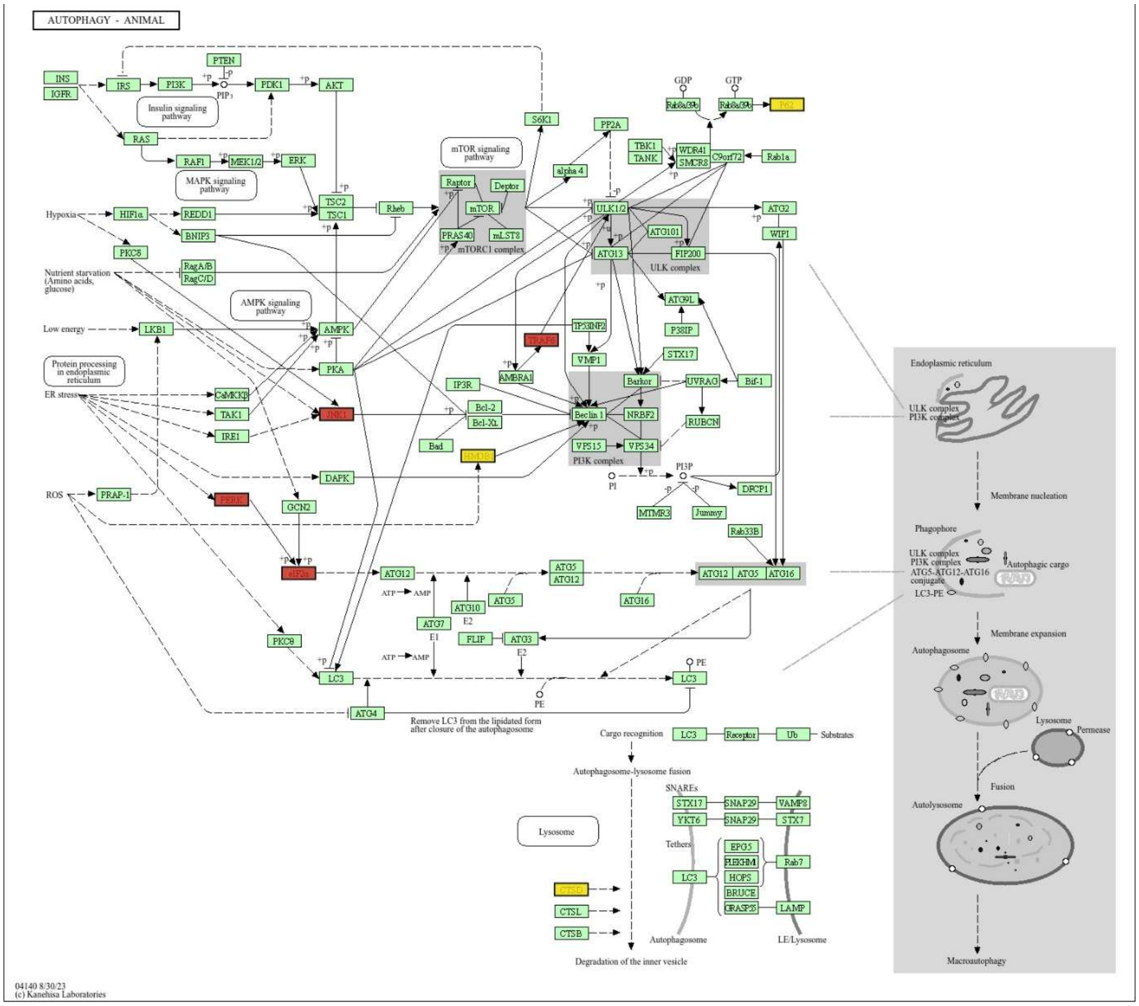


Supplementary Figure 9. NEPTUNE highlighted direct markers of autophagy across all timepoints in the cones of *rd1* mice. Red labels indicate downregulated markers while yellow labels indicate upregulated markers. Pale colours show weak regulation dictated by the |log2FC| values. Green colour indicates all the other genes of the pathway.

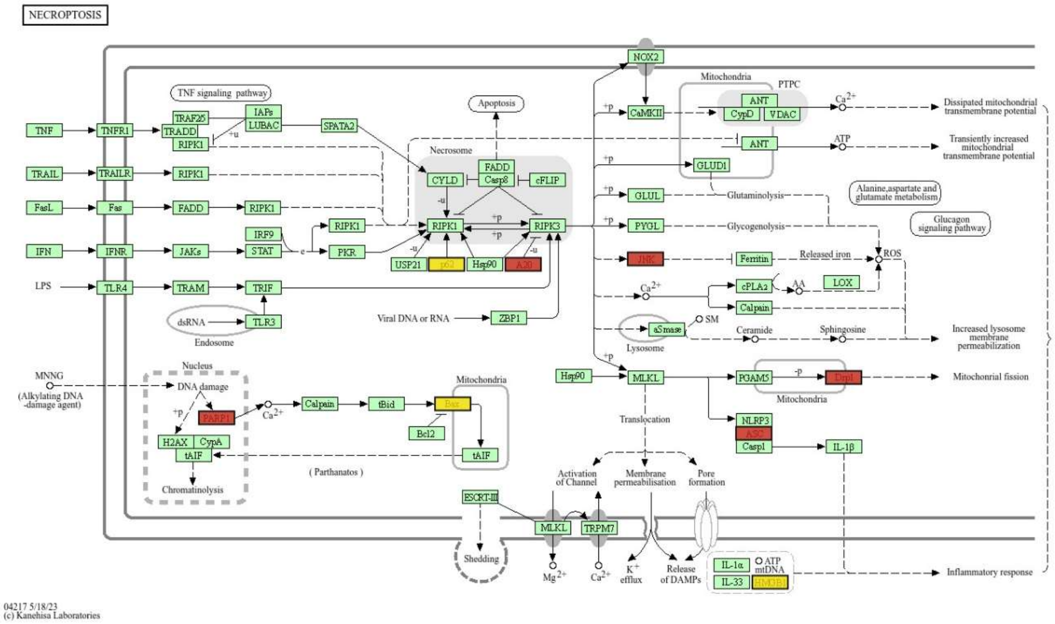


Supplementary Figure 10. NEPTUNE highlighted direct markers of necroptosis across all timepoints *rd1* cones. Red labels indicate downregulated markers while yellow labels indicate upregulated markers. Pale colours show weak regulation dictated by the |log2FC| values. Green colour indicates all the other genes of the pathway.

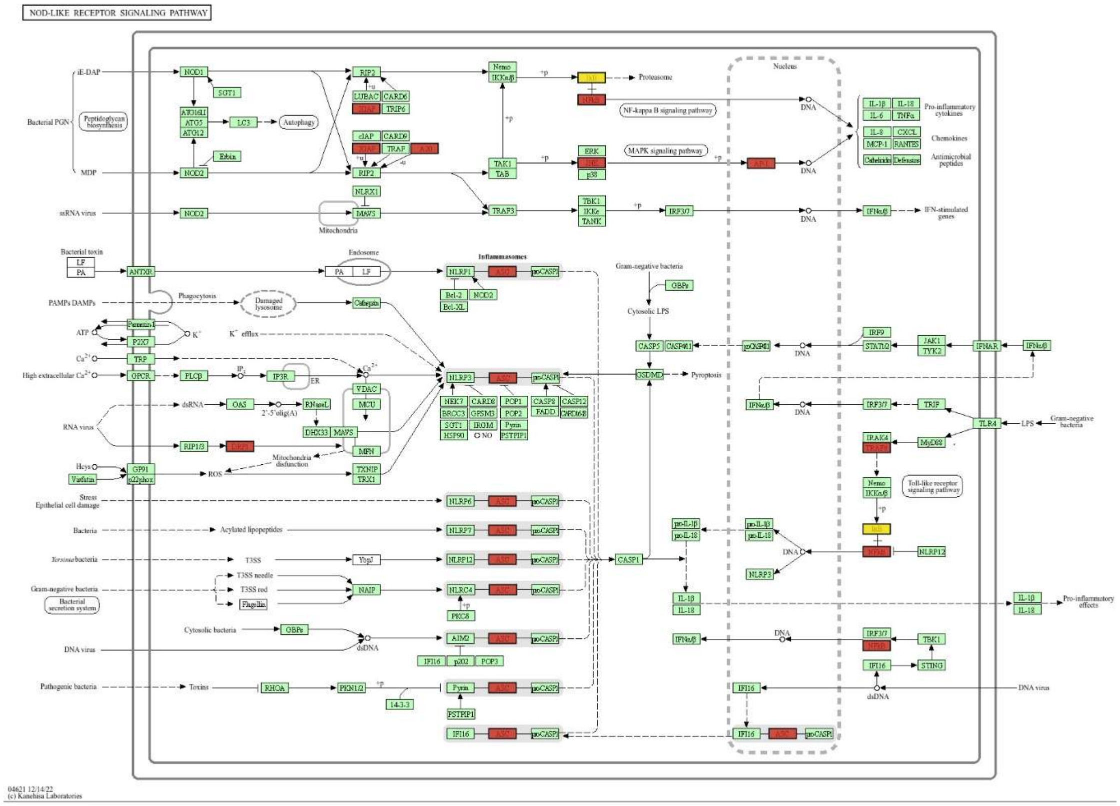


Supplementary Figure 11. NEPTUNE highlighted direct markers of pyroptosis across all timepoints in the cones of *rd1* mice. Red labels indicate downregulated markers while yellow labels indicate upregulated markers. Pale colors show weak regulation dictated by the |log2FC| values. Green colour indicates all the other genes of the pathway.

Supplementary Table 1. Cell counts of rods and cones.

| | Rods | Cones |
|---|---|---|
| WT_P11 | 5204 | 92 |
| WT_P13 | 7924 | 172 |
| WT_P17 | 6319 | 111 |
| rd1_P11 | 5627 | 221 |
| rd1_P13 | 5803 | 495 |
| rd1_P17 | 917 | 653 |

*Note: Numbers in the table mentions the cell counts.*
